# Early Events in G-quadruplex Folding Captured by Time-Resolved Small-Angle X-Ray Scattering

**DOI:** 10.1101/2024.09.05.611539

**Authors:** Robert C. Monsen, T. Michael Sabo, Robert Gray, Jesse B. Hopkins, Jonathan B. Chaires

## Abstract

Time-resolved small-angle X-ray experiments (TR-SAXS) are reported here that capture and quantify a previously unknown rapid collapse of the unfolded oligonucleotide as an early step in G4 folding of hybrid 1 and hybrid 2 telomeric G-quadruplex structures. The rapid collapse, initiated by a pH jump, is characterized by an exponential decrease in the radius of gyration from 20.6 to 12.6 Å. The collapse is monophasic and is complete in less than 600 ms. Additional hand-mixing pH-jump kinetic studies show that slower kinetic steps follow the collapse. The folded and unfolded states at equilibrium were further characterized by SAXS studies and other biophysical tools, to show that G4 unfolding was complete at alkaline pH, but not in LiCl solution as is often claimed. The SAXS Ensemble Optimization Method (EOM) analysis reveals models of the unfolded state as a dynamic ensemble of flexible oligonucleotide chains with a variety of transient hairpin structures. These results suggest a G4 folding pathway in which a rapid collapse, analogous to molten globule formation seen in proteins, is followed by a confined conformational search within the collapsed particle to form the native contacts ultimately found in the stable folded form.

## Introduction

Understanding the folding mechanism of biological molecules is fundamental to understanding their form and function. G-quadruplexes (G4) are unique DNA or RNA structures whose folding, not surprisingly, differs in detail from the folding of proteins, RNA, or duplex DNA. G4 dynamics and the diversity of its folding mechanisms were recently reviewed^[1]^. The folding of G4 structures formed by human telomeric repeat sequences has been studied in the most detail by a variety of experimental approaches, including absorbance, fluorescence and circular dichroism spectroscopies, NMR, mass spectrometry, single-molecule methods and molecular dynamics simulations^[2]^. No consensus on the detailed folding mechanism has emerged, apart from the fact that it is a multistep, slow process. Folding funnels, kinetics partitioning and sequential folding models with populated intermediates were suggested by different investigators. Many specific questions about the process remain. Because of limitations in the time resolution of most of the methods used, early steps in the folding of telomeric sequences are poorly characterized. Stopped-flow absorbance studies inferred from the cation dependency of the fastest measurable relaxation time that a kinetic process faster than the 20ms deadtime of the instrument existed^[2c]^. Stopped-flow absorbance, CD, and FRET studies from our laboratory^[2c, 2h]^ captured early cation-driven folding events that were complete within 1-2 seconds. The FRET results indicated a fast reduction in the end-to-end distance of oligonucleotide chain, but no quantitative estimate of that distance was possible. A time-resolved CD spectrum was obtained by stopped-flow^[2h]^ of an intermediate species formed at 1 s with a spectrum characteristic of an antiparallel basket form^[3]^(or perhaps a two-tetrad basket antiparallel structure^[4]^).

More recent microfluidic mixing^[5]^ and T-jump^[6]^ methods, with micro- to millisecond resolution, used FRET or CD detection to detect faster events. The mixing experiments were initiated by cation concentration jumps, and the fast events (20 -40 µs lifetimes) observed were interpreted as the formation of hairpin loops. T-jump experiments monitored perturbations of the folded telomeric “basket” G4 structure in Na^+^ solution and monophasic fast relaxations in the ms to s range were observed over a range of cation concentrations and temperatures. The underlying molecular origin of these relaxations was not clear although it was noted that they were much slower than expected for simple hairpin formation. A new type of micromixer with a deadtime of 40 µs and a wide time window allowing continuous data collection up to 300 ms detected multiple relaxation times by FRET for the folding the human telomere in KCL or NaCl^[7]^. The FRET signal was not converted to quantitative distance information. The interpretation of the multiple relaxation times as a sequential mechanism that includes a triplex intermediate is flawed by the failure of the authors to recognize that such a species forms only on a much slower time scale^[2h]^ than was measured using the new micromixer and that complete folding requires much longer than 300 ms^[2c, 2h, 2i, 2k]^.

A full description of G4 folding remains beyond the reach of current molecular dynamics simulation methods. MD simulation can at best reach a ms timescale, well below the actual experimentally observed timescale of the folding process. MD results suggest that the folding process may be an extreme case of kinetic partitioning, and that the full G-quadruplex folding landscape is not reducible to a few order parameters or measurables. A number of pre-folded structures, such as hairpins and triplexes, were calculated to be stable, and G-quadruplexes with strand-slippage were proposed as potential off-pathway intermediates^[2m]^. Using NMR methods, Frelih and colleagues showed that, indeed, both strand-slipped triplex intermediates and 5’ hairpin features are observed to form from the telomere sequences under low temperature, low salt, and low pH conditions^[8]^.

We report here the results of rapid-mixing, time-resolved small-angle X-ray scattering (TR-SAXS) experiments intended to better characterize the early events in G4 folding. The folding of both the human telomere hybrid 1, 5’TTGGG(TTAGGG<u>)_3_</u>A, (“2GKU”) and hybrid 2, 5’TAGGG(TTAGGG<u>)_3_</u>TT, (“2JSL”) structures in K+ solution was studied. A continuous laminar-flow TR-SAXS apparatus with millisecond resolution^[9]^ was used to affect a pH jump to initiate G4 folding, with scattering profiles then collected at intervals over the range 1 ms – 1.2 s. The significant advantage of this TR-SAXS approach is that it allows rapid changes in a well-defined structural parameter, the radius of gyration (R_g_), to be monitored. To the best of our knowledge, this is the first use of time-resolved SAXS to characterize G4 folding.

## Results and Discussion

### Equilibrium conformational analysis of the folded, unfolded, and Li^+^ destabilized states

To understand the telomere G4 folding pathway, we first identified conditions that favor the folded or fully unfolded forms. Unfortunately, chemical denaturants like urea and guanidine-HCL commonly used to unfold proteins or RNA do not unfold G4 structures^[10]^. G-quadruplexes are stabilized by a number of factors, including loop length, crowding effects, pH, temperature, and ionic strength and composition. G4 formation is mainly driven by hydrogen bonding (Hoogsteen bonding of guanines in tetrads and loop “capping” structures that span either terminus), electrostatics (ion coordination in both the central G-tetrad channel and along the negatively charged backbone) and nucleotide stacking (pi-stacking of gaunines and van Der waals stabilization). The particular conformation of a G4 strongly depends on the identity of the central channel cation. For instance, the human telomere G4 sequence 143D adopts an antiparallel conformation in the presence of Na^+^ cations in solution^[11]^, a parallel conformation in the presence of K^+^ cations in dehydrated, crowded crystallization conditions^[12]^, and an equilibrium of hybrid-1 and hybrid-2 conformations in the presence of K^+^ cations in solution^[13]^. Other cations, such as Li^+^, have been reported as having destabilizing effects on G4 stability^[14]^, with Li^+^ often used in place of K^+^ and Na^+^ ions in hopes of preventing G4 formation^[8]^.

**Figures 1, S1, and 2** present the results of equilibrium studies characterizing the folded form of 2GKU in KCl solution compared to its alkaline and thermally denatured forms, to a reference single-strand oligonucleotide (dT_24_), and to 2GKU in LiCl solution. The CD spectra in **figure 1A** show the characteristic spectrum of the hybrid 1 folded form for 2GKU in KCl^[3]^, in contrast to the nearly flat, featureless spectra for the alkaline and thermally denatured forms. Replacing K^+^ ions with Li^+^ (blue curve) results in a spectrum with a near zero magnitude at 240 nm, but with significant signal in the range of 260-300 nm, indicating that nucleotide stacking interactions persist^[15]^. The observed spectrum is inconsistent with any reported G4 fold^[3]^. **Figure S1** shows the matched ^1^H-NMR spectra for each condition. Only in KCl solution does 2GKU show the characteristic 12 G-tetrad resonances at 10-12 ppm expected for a folded, hybrid 1 G4. Under alkaline denaturation conditions, no secondary structure proton shifts are observed in the ∼8-14.5 ppm range. In Li^+^ buffer at pH 7.2, 2GKU is not entirely denatured, as evident by the weak signals observed in the G-tetrad imino fingerprint region (10-12 ppm), some of which overlap with the chemical shifts of the folded hybrid-1 conformation. The same is observed for 2JSL under identical conditions (**Figure S2**).

**Figure 1.**
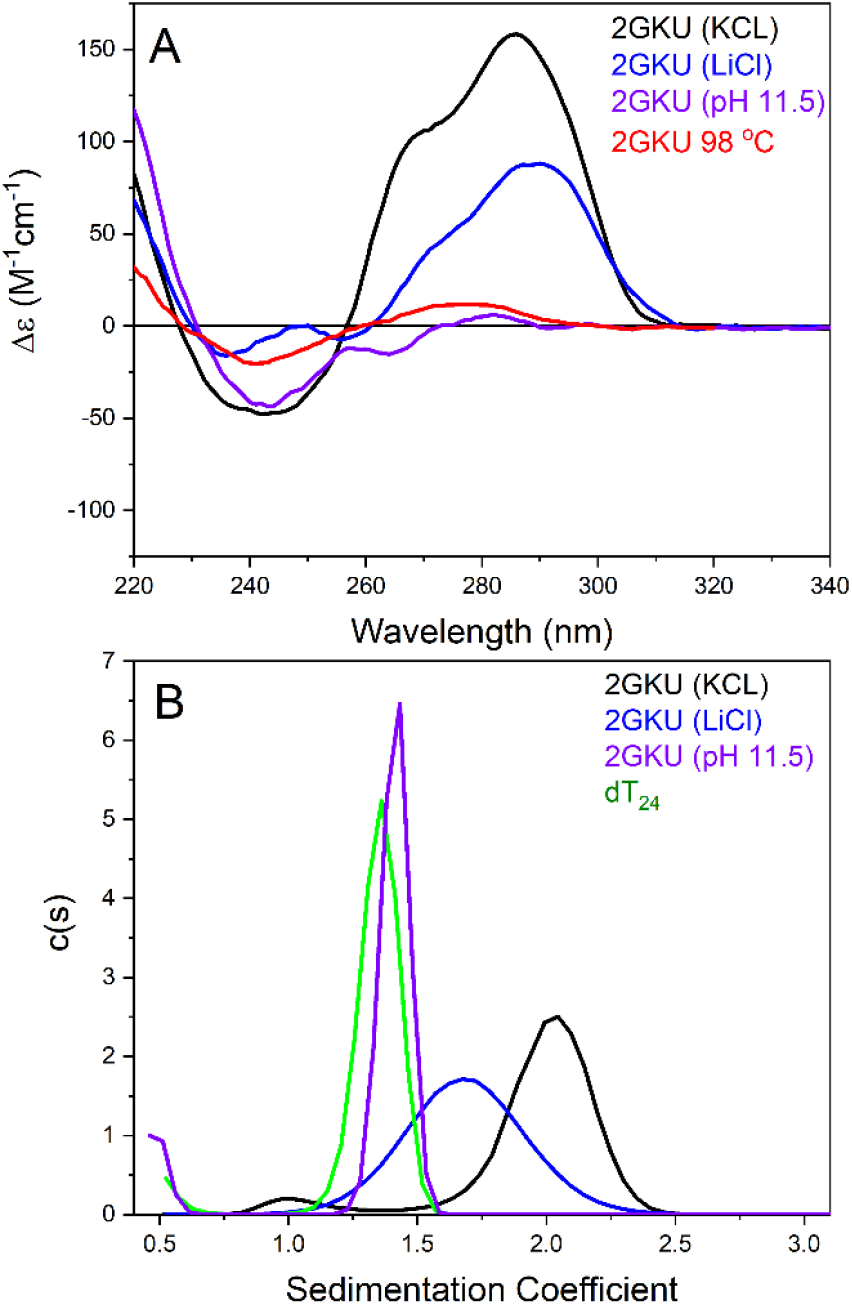
(A) CD spectra of 2GKU in KCL (black), LiCl (blue), KCl at pH 11.5 (purple) and KCl at 98 °C (red). (B) Sedimentation coefficient distributions for 2GKU in KCl (black), LiCl (blue) and KCl at pH 11.5 (purple). The sedimentation of an unstructured single-stranded reference oligonucleotide, T_24_, is shown in green.

Although heat denaturation may be effective for ensuring a fully unfolded G4 sequence, temperature jump kinetic experiments are difficult in practice, and temperature directly affects molecular diffusion, complicating folding analysis^[16]^. Also, orthogonal methods like SAXS, AUC-SV, and NMR are difficult at the high temperatures required to melt G4 DNA. Therefore, alkaline denaturation was chosen for subsequent folding analyses. The use of alkaline denaturation in kinetics studies of duplex DNA formation has a long history^[17]^.

**Figure 1B** shows the results of AUC-SV analysis of the 2GKU sequence in K^+^ buffer at pH 7.2 and 11.5, Li^+^ buffer at pH 7.2, and an oligonucleotide of similar length to 2GKU consisting of only thymine, d(T)_24_, which was also in K^+^ buffer at pH 7.2. The fully folded hybrid-1 form of 2GKU (black line) sediments with S_20,w_ = 2.0, whereas at pH 11.5 it sediments at nearly the same rate as the d(T)_24_ sequence (1.6 S_20,w_ vs. 1.4 S_20,w_), indicating that it is unfolded. In the presence of Li^+^, 2GKU sediments at a rate intermediate to the fully unfolded and folded extremes (1.8 S_20,w_), indicating that it is more extended than the folded 2GKU, yet more compact than the pH denatured 2GKU, possibly existing as a fast equilibrium between the compact and extended states. The same trend is observed for 2JSL (**Figure S3**).

We next evaluated the global structural features of 2GKU in the same conditions using equilibrium SAXS. **Figure 2A-C** shows the background subtracted scattering, dimensionless Kratky, and P(r) distribution plots, respectively, for 2GKU at pH 7.2 with KCl (black), LiCl (red), and pH 11.5 with KCl (blue) (**Figures S4-9** show each SEC-SAXS elution profile). Under alkaline conditions, 2GKU is maximally extended and flexible, given by the elevated plateau at high qR_g_ and right skew to more than double the D_max_ of the folded 2GKU and nearly identical to poly dT_24_ under identical conditions (**Figure S10**). The measured D_max_ is larger than the end-to-end distance of a rigid single-stranded sequence with the helicity found in a B-form conformation (experimental D_max,exp._ = ∼83 Å vs B-form calculated D_max,calc._ = 75 Å). Conversely, at pH 7.2 in the presence of either counterion, 2GKU is very compact and globular, given by the near symmetric bell and Gaussian shapes of the distributions, respectively. We have previously compared the K^+^-solution NMR structure of 2GKU to its equilibrium scattering and confirmed an excellent fit to the SAXS data^[18]^. An increase in D_max_ of 5 Å and 12 Å was observed for the LiCl condition for 2GKU and 2JSL (**Table 1, Figure 2 and S11**), respectively, in agreement with the increases in frictional coefficient (*f/f*_*o*_) observed in AUC-SV measurements.

**Table 1.** Physical properties of 2GKU and 2JSL under different solution conditions.

| Sequence | 2GKU | 2GKU | 2GKU | 2JSL | 2JSL | 2JSL |
| --- | --- | --- | --- | --- | --- | --- |
| PROPERTY | KCL | LiCl | KCL | KCL | LiCl | KCL |
|  | PH 7.2 | PH 7.2 | PH | PH 7.2 | PH 7.2 | PH |
|  |  |  | 11.5 |  |  | 11.5 |
| $S_{20,W}$ | 2.02 | 1.76 | 1.62 | 2.05 | 1.72 | 1.64 |
| $F/F_0$ | 1.37 | 1.52 | 1.79 | 1.41 | 1.51 | 1.81 |
| $R_s$, Å | 16.9 | 18.3 | 22.5 | 17.8 | 28.2 | 23.2 |
| $R_g$, Å | 12.27 | 11.92 | 24.26 | 12.13 | 12.89 | 24.26 |
| (GUINIER) | $\pm 0.04$ | $\pm 0.01$ | $\pm 0.17$ | $\pm 0.03$ | $\pm 0.04$ | $\pm 0.11$ |
| $R_g$, Å | 12.23 | 11.97 | 24.56 | 12.2 | 13.18 | 25.03 |
| (GNOM) | $\pm 0.03$ | $\pm 0.02$ | $\pm 0.11$ | $\pm 0.03$ | $\pm 0.10$ | $\pm 0.08$ |
| $D_{MAX}$, Å | 38 | 43 | 83 | 39 | 51 | 87 |

**Figure 2.**
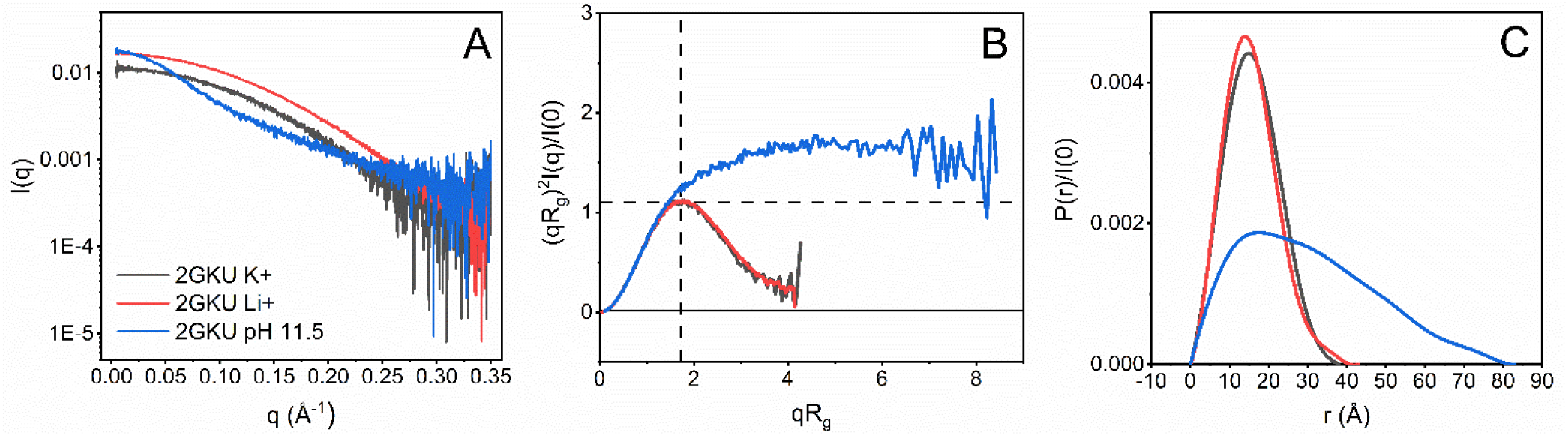
Equilibrium SAXS analysis of 2GKU. (A) Scattering profiles, (B) Dimensionless (re-binned) Kratky plots, and (C) Normalized P(r) distributions of 2GKU in KCl at pH 7.2 (black), LiCl at pH 7.2 (red), and KCl at pH 11.5 (blue).

The exact nature of the denatured state is important for understanding the folding process, but little is known about it for G4 forming sequences. SAXS offers a unique capability for understanding its properties, as shown in **Figure 3**. SAXS scattering curves are a weighted composite of all solution scatterers present. Therefore, flexibility as seen in **Figure 2** (blue curve) should be interpreted as an ensemble of flexible structures. To this end, we used the Ensemble Optimization Method (EOM)^[19]^ to determine a set of conformers from implicit solvent molecular dynamics simulations that have theoretical scattering curves that recapitulate the experimental data. **Figure 3A-B** shows the EOM results as R_g_ and D_max_ distributions of the conformer pool (black lines) and selected ensemble (red lines), respectively. **Figure 3C** shows the scattering profile of the alkaline denatured 2GKU with an EOM ensemble fit overlaid (red) and with fit residuals shown below. The χ^2^-value of 1.162 and normally distributed residuals indicate a satisfactory fit of the ensemble of conformers shown in **Figure 3D**. Moreover, the resulting representative ensembles have R_g_ values that are in excellent agreement with what was measured (2GKU: R_g,ensemble_ = 24.56 Å vs. R_g,experimental_ = 24.56 ± 0.11 Å and 2JSL: R_g,ensemble_ = 25.54 Å vs. R_g,experimental_ = 25.03 ± 0.08 Å). Representative 2GKU models in **Figure 3D** are displayed from most to least extended going from top left to bottom right. The D_max_ range for the conformers is 59 – 89 Å and the fractional makeup is given in the figure for each model. Interestingly, most of the models constituting the EOM ensembles (∼92% weight) have guanine hairpin features that formed transiently throughout the simulations. 2JSL had a similar result, albeit with 100% of the ensemble having 5’ hairpin features included (**Figure S12**).

**Figure 3.**
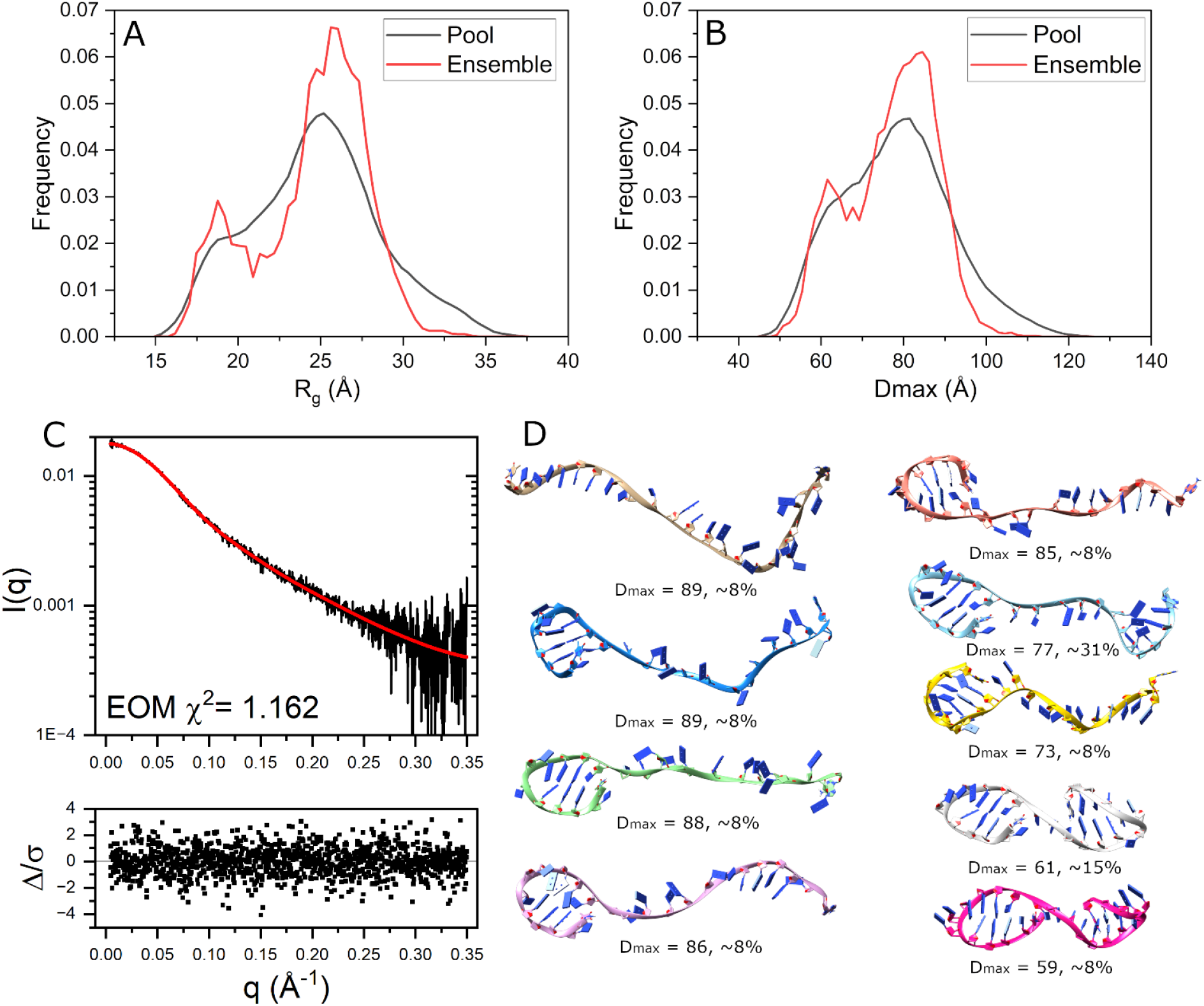
EOM analysis results for 2GKU. (A-B) EOM distributions for the radius of gyration (A) and D_max_ (B) for the total pool of conformers (black) and the selected ensemble of flexible structures (red). (C) pH 11.5 scattering curve with EOM fit overlaid in red and residuals below. (D) Best fit ensemble of conformers chosen by EOM from duplicate 500 ns implicitly solvated MD simulations starting from single-stranded 2GKU showing the most extended (top) to the most compact conformations (bottom) oriented with 5’ end on the left. Calculated D_max_ values and % ensemble weight are given above each model. EOM statistics: R_flex_ (random) / R_sigma_: ∼ 84.25% (∼ 90.67%) / 0.85.

### TR-SAXS reveals millisecond timescale collapse to a pre-folded intermediate

Hand-mixing pH-jump experiments monitored by CD were done to ensure that a rapid adjustment from alkaline to a neutral pH gave folding kinetics that were consistent with cation induced folding. **Figure 4A** shows the CD spectral changes of the 2GKU alkaline to neutral pH jump monitored over 11,000 seconds with a mixing dead time of ∼30 seconds. Analyzing the data using a singular value decomposition (SVD) reveals that over this time frame two intermediates are formed with relaxation times of approximately 15 minutes and 1.5 hours, followed by a much longer relaxation to the final folded form of about 10 hours. The final spectrum is in qualitative agreement with what is expected for the folded hybrid 1. Upon the jump to neutral pH, component 1 (red) immediately forms, and exhibits peaks at 250 and 290 nm that are consistent with folding intermediates that have been previously observed by K^+^-induced folding of the same sequence^[2h]^ and a mass spectrometry kinetic study^[2k]^ that proposed a two-tetrad antiparallel basket conformation as an intermediate^[4b]^. The previous kinetic studies investigating the K^+^-induced 2GKU folding at the millisecond timescale show that the earliest spectral intermediate forms within about 300 milliseconds and exhibits a positive 290 nm peak and negative ellipticity at 260 nm^[2h]^, consistent with an antiparallel fold^[20]^. However, similar hand-mixing pH jump NMR experiments, deconvoluted with SVD, show complex imino shifts that are not easily assigned to a single G4 fold (**Figure S13**).

**Figure 4.**
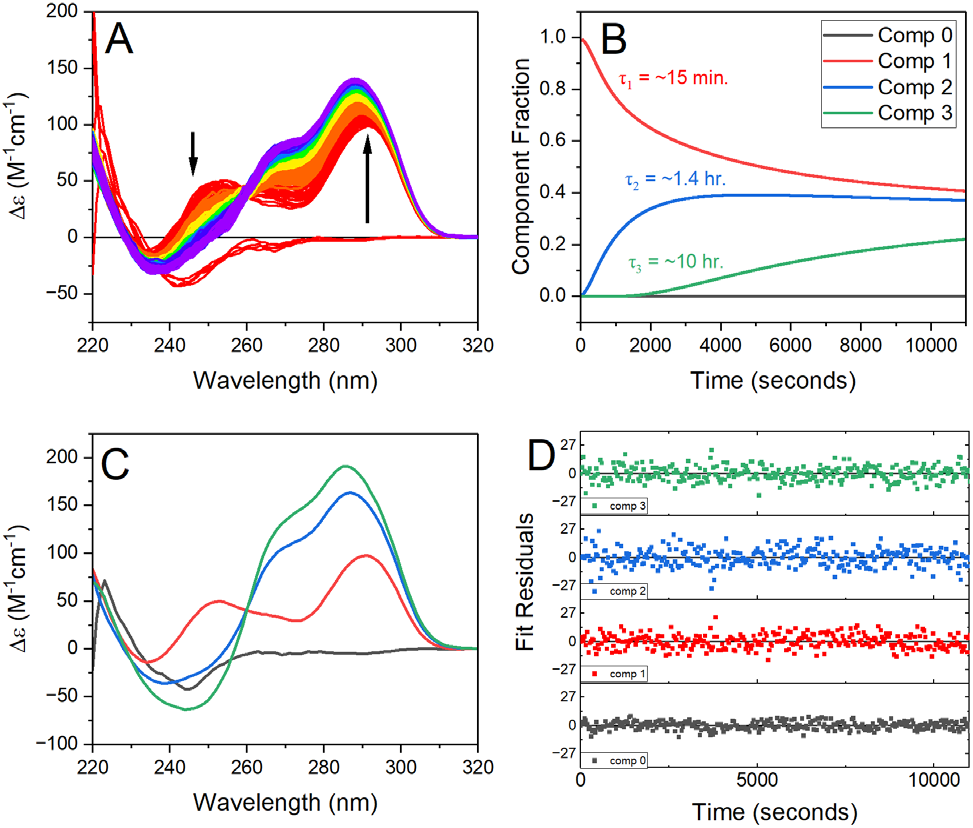
2GKU pH-jump spectra and kinetics. Folding was initiated by manually mixing a small volume of concentrated hydrochloric acid into the alkaline (pH 11.5) buffer solution to drop it to neutral pH (final pH = 7.1). (A) CD spectra of the alkaline denatured 2GKU pre- and post-pH-jump monitored over 11,000 s. (B) Fractional component plot from SVD with relaxation times overlaid based on single exponential fits. (C) Deconvoluted CD spectra for each component in B from SVD. (D) Residuals from the nonlinear least-squares fit to the data from SVD analysis.

Time-resolved SAXS experiments to provide insight into the structural collapse following imino protonation on a sub-second time scale. **Figures 5A-B** show the normalized P(r) distribution and dimensionless Kratky plots derived from the scattering time series for the pH-induced folding of 2GKU monitored over 1-1,200 ms. For clarity, the Kratky distributions have been re-binned, and only select time points shown. From these plots it is evident that between time 0 s (i.e., the pH 11.5 equilibrium SAXS data) and ∼1 ms there is a noticeable reduction in D_max_ values commensurate with changes in the Kratky region of 3.6 - 8 qR_g_, suggesting that a sub-millisecond event occurs prior to the start of our TR-SAXS measurements. DNA hairpin formation occurs on timescales of microseconds or less^[21]^, and recent TeZla micromixer fluorescence experiments of the FRET-labeled 143D telomere G4, (using K^+^-induced folding) have shown that the telomere hairpins appear in under 100 µs^[7]^. Those results suggest that our measurements may have missed rapid hairpin formation event although our equilibrium SAXS modeling with EOM clearly suggested their presence. Inspection of curves from the final time points (861 and 1205 ms) indicates that the collapse from the hairpin ensemble to globular pre-folded intermediate is complete well within 1 second. Direct fitting of the derived R_g_ values *versus* time using a single exponential decay model reveals a relaxation rate of τ = 187 ± 13 ms (**Figure S14**) that is in agreement with values obtained by prior K^+^-induced folding measured by stopped-flow absorbance (τ_285 nm_ = 104 ± 4 ms, τ_265 nm_ = 260 ± 20 ms)^[2h]^.

**Figure 5.**
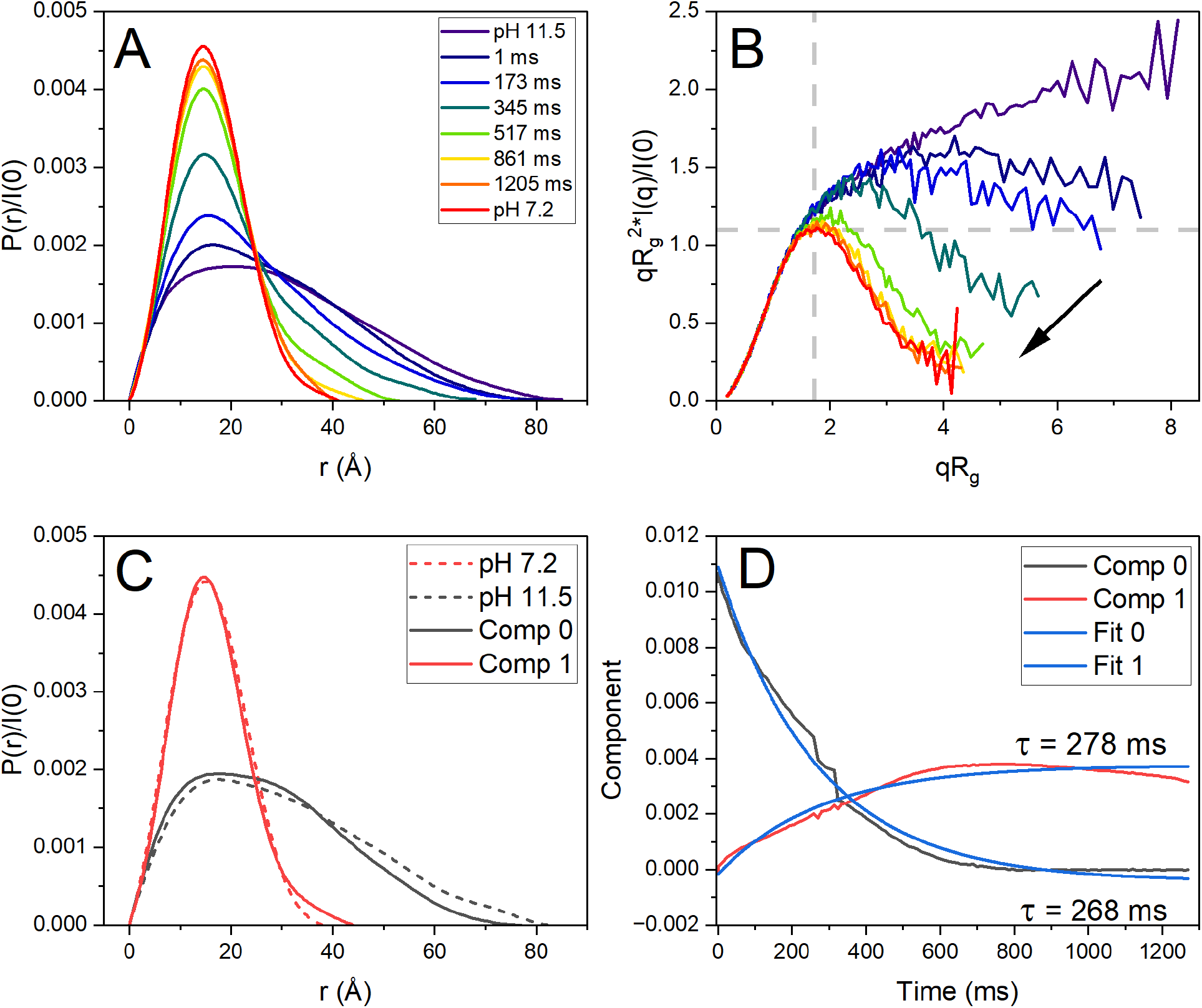
Time-resolved SAXS results of 2GKU following pH jump from 11.5 to 7.2. (A) Selected normalized P(r) distributions of the pH-induced structural collapse of 2GKU bracketed by the equilibrium SAXS profiles showing the conversion from an extended unstructured species to a globular and compact particle of nearly identical size and shape as the folded hybrid 1 form. (B) Dimensionless Kratky plots (data re-binned for clarity with log mode and re-bin factor 4) showing the transition from the denatured flexible chain to a compact globular form that is nearly identical to the equilibrium 2GKU scattering. (C) REGALS derived regularized P(r) distributions comparing the two deconvoluted components (solid lines) with the equilibrium SAXS 2GKU distributions (dashed lines). (D) Regularized component concentration profiles from REGALS deconvolution with single exponential decay relaxation times overlaid and fits shown in blue. Red and black curves correspond to the solid red and black P(r) distributions in C.

As the TR-SAXS kinetic data set is complex and could contain unanticipated evolving species, we employed REGularized Alternating Least Squares (REGALS)^[22]^ analysis within the BioXTAS RAW^[23]^ package to deconvolve the time-resolved scattering data. In both 2GKU and 2JSL data sets only two components were identified from SVD, corresponding to the pre- and post-collapsed folding intermediates. REGALS was then run with our D_max_ values from equilibrium SAXS as estimated constraints to obtain the deconvolved spectra shown in **Figure 5C** with corresponding component concentration profiles in **Figure 5D**. The resulting P(r) distributions are physically reasonable and in good agreement with the equilibrium P(r) distributions (dashed lines). Fitting the component concentration curves in **Figure 5D** with single exponential functions yields relaxation times of 278 and 268 ms for the formation of the collapsed structure and the disappearance of the pre-folded structure, respectively. 2JSL also had only two components and the REGALS deconvolution showed a collapse with similar relaxation times of ∼290 ms (**Figure S15**). Again, these rates of collapse are in excellent agreement with previous spectroscopic stopped flow investigations^[2h]^ and the recent TeZla mixing device results using the FRET-labeled 143D^[7]^.

The power of time-resolved studies by SAXS (and analysis with REGALS) is that along with the kinetic analyses and qualitative descriptions, there are also quantitative structural properties that aid in describing the nature of the pre- and post-collapse species. **Table 2** compares the equilibrium and time-resolved SAXS R_g_ and D_max_ values for 2GKU and 2JSL. As discussed in the description of the P(r) and Kratky data above, the D_max_ and R_g_ values of the equilibrium alkaline denatured species are much larger than the earliest time points of the TR-SAXS measurements, which suggests sub-millisecond events are occurring, such as a shift to all hairpin structures. Similarly, although the collapse has finished by our latest time points, the D_max_ and R_g_ values of the TR-SAXS species are larger than for the fully folded and equilibrated 2GKU and 2JSL, indicating that their folding is not complete.

**Table 2.** Comparison of equilibrium (EQ) and time-resolved (TR-SAXS) results of 2GKU and 2JSL comparing the equilibrium denatured and folded results in KCl to the initial and final species from time-resolved SAXS. See also **Table S1**.

| Sequence | 2GKU | 2GKU | 2GKU | 2GKU | 2JSL | 2JSL | 2JSL | 2JSL |
| --- | --- | --- | --- | --- | --- | --- | --- | --- |
| Property | EQ-SAXS<br>(KCl) | TR-SAXS<br>(final) | EQ-SAXS<br>(KCl pH 11.5) | TR-SAXS<br>(initial) | EQ-SAXS<br>(KCl) | TR-SAXS<br>(final) | EQ-SAXS<br>(KCl pH 11.5) | TR-SAXS<br>(initial) |
| $R_g$, Å<br>(GUINIER) | $12.27 \pm 0.04$ | $12.63 \pm 0.01$ | $24.26 \pm 0.17$ | $20.65 \pm 0.03$ | $12.13 \pm 0.03$ | $12.80 \pm 0.01$ | $24.26 \pm 0.11$ | $21.57 \pm 0.05$ |
| $R_g$, Å (GNOM) | $12.23 \pm 0.03$ | $12.84 \pm 0.02$ | $24.56 \pm 0.11$ | $21.63 \pm 0.04$ | $12.20 \pm 0.03$ | $13.02 \pm 0.03$ | $25.03 \pm 0.08$ | $22.40 \pm 0.05$ |
| $D_{\text{max}}$, Å | 38 | 48 | 83 | 73 | 39 | 48 | 87 | 79 |

## Conclusion

These data show that an early step in G4 folding, complete in less than 1 s, is the collapse from an extended, flexible, polynucleotide chain to a compact form. For the 2GKU sequence, the radius of gyration decreases from 21.6 to 12.8 Å, while the maximal interparticle distance decreases from 83 to 48 Å. The collapsed particle is slightly larger than the fully folded form at equilibrium (**Table 2**). These quantitative measures provide structural details that must be accounted for in any kinetic or computational model of the folding process. Kinetic steps both precede and follow this collapse. Events faster than the 1 ms deadtime of the TR-SAXS apparatus may plausibly be attributed to transient hairpin formation, especially since the scattering curve for the unfolded state at equilibrium is best modeled as an ensemble of largely single-stranded forms with a variety of transient hairpins (**Figure 3**). Kinetic studies with µs time resolutions have proposed hairpin formation as an initial step in G4 folding^[5, 7]^. Following the rapid collapse, additional kinetic steps occur over 1-10000 s that can be observed by UV absorbance, CD, FRET, NMR and mass spectrometry^[2c, 2f, 2h, 2i, 2k]^. These studies show that long-lived intermediates exist, since the observation of characteristic spectral or charge states requires that species accumulate at sufficient concentrations to produce a measurable signal. Specific cation coordination to stabilize the final folded G4 conformation seems to occur after the collapse reported here ^[2k]^.

The conformational freedom of the unfolded polynucleotide chain provides a favorable entropy contribution that opposes G4 folding and stabilizes the denatured state. With rotation around 6 backbone torsion angles for each nucleotide, along with rotation around the glycosidic bond that attaches the base, an estimate for the Boltzmann conformational entropy of the 2GKU sequence (with n=24 nt) is S= Rl*n*(7^n-1^)= R*ln*(7^23^)= 88.6 cal/K-mol, where R is the gas constant and the exponential term enumerates the number of possible microstates. Experimental calorimetric measurements^[24]^ of 2GKU unfolding determined a favorable entropy of 60.4 cal/K-mol for the first step in a sequential folding process, in good agreement with the theoretical estimate given that there are surely other coupled contributions within the experimental value. These entropy estimates suggest that the loss of conformational freedom presents a free energy barrier (ΔG_conf_= -TΔS) of 18 - 26 kcal/mol that must be overcome for folding. The favorable free energy of folding arises from favorable contributions from molecular interactions such as hydrogen bonding, base stacking, and cation coordination. But an often overlooked contribution from the hydrophobic force resulting from the removal of solvent accessible surface area also drives folding. The cited experimental calorimetric study^[24]^ accurately determined the heat capacity change for the first step in the 2GKU folding process to be -420 cal/K-mol. Spolar and Record^[25]^ determined an empirical relationship between the hydrophobic free energy contribution (ΔG_hyd_) and the heat capacity change (ΔC_p_), ΔG_hyd_ = 80 ΔC_p._. From this relation, a free energy contribution of -33.6 kcal/mol from the hydrophobic force overcomes the entropic barrier and drives the first step in the folding of 2GKU. The origin of this contribution is the removal of solvent accessible surface area (SASA) with a concomitant release of water. Hadzi and Lah^[26]^ determined the correlation ΔC_p_= 0.134(ΔSASA), where ΔSASA is the change in SASA, from which an estimate of the removal of about 3100 Å^2^ of surface area results for 2GKU folding. Using the model weights from the EOM ensemble to more accurately model the SASA of the unfolded state (**Figure 3**), we can approximate the weighted change in SASA for 2GKU’s collapse to be 2477 Å^2^. This estimate results in a predicted heat capacity change of ΔC_p_= 0.134(−2477 Å^2^) = -332 cal/K-mol, which is close to the experimental measured value (i.e., -420 cal/K-mol), and ΔG_hyd_ = -26.6 kcal/mol, which is approximately the maximum estimated energy needed to overcome the entropic cost of folding. Our TR-SAXS measurements have, for the first time, provided the structural details and basis of this initial hydrophobic collapse.

Rather than offering a formal kinetic mechanism for the overall folding process, it is perhaps more useful to provide a phenomenological description of events that occur that must be accounted for in any mechanistic proposal. Below one millisecond, transient hairpin formation likely occurs that have been experimentally observed^[27]^ and captured by our EOM model (**Figure 3**). MD simulations seem to capture many of the bewildering ensemble of possible hairpins that might form in this phase^[2m]^. A stable hairpin structure was detected by NMR in the 5’ end of the human telomer G4 sequence^[8]^. At about 1 ms, the hydrophobic collapse described here to a “molten globule”-type structure occurs. The collapse is complete by 1 s. FRET changes occur on this same timescale indicating that the polynucleotide end-to-end distance is drastically decreased^[2h]^. SAXS provides a quantitative description of the global structural features of this collapsed structure that could be used to test molecular simulations of the process (**Table 2**). Next, a time-resolved CD spectrum shows that by 2 s an intermediate has formed with the spectral characteristics of an antiparallel G4 structure^[2h]^. Over the 5-1000 s timescale, FRET and CD changes are stable^[2h]^, and mass spectrometry (indicative of cation binding) shows the transient formation of a +1 charge species that transforms into a +2 charge species^[2k]^. By 1000 s, the +1 charge species has disappeared, leaving only the +2 species. During this phase, there are complex, position-dependent, changes in the fluorescence of the site-specific probe 2-aminopurine substituted at specific locations in the G4 sequence, indicative of strand rearrangement^[2h]^. The development of NMR spectra characteristic of G4 hybrid structures begins around 100 s (**Figure S13**), perhaps earlier but inaccessible because of the deadtime of NMR mixing experiments used^[2i]^. From 1000 -10000 s, NMR spectra evolve to a spectrum characteristic of a G4 hybrid 1 spectrum (like 2GKU). During this phase, subtle CD and 2-aminopurine fluorescence changes occur until equilibrium is reached^[2h]^. Overall, during these phases discrete populated intermediates come and go, leading to the formation of G4 hybrid structures at the final equilibrium. Discrete intermediates must form in sufficient concentrations to provide the biophysical signals observed over these phases.

As an overall global description of the folding process, we propose that the unfolded polynucleotide chain is in fact never fully extended but instead is an ensemble of transient hairpin structures that collapses into a “molten globule” structure at the onset of folding, driven primarily by hydrophobic forces. “Kinetic partitioning” may well describe this collapse. Within the collapsed particle, a conformational selection process may then occur within the confined space of the globule, with specific cation binding and coordination serving to stabilize on-pathway intermediates and the final native conformation. In this view, G4 folding is neither a kinetic partitioning phenomenon nor discrete sequential folding pathway - it is both.

## Supporting Information

The authors have cited additional references within the Supporting Information.^[3, 19, 22-23, 28]^

## Acknowledgements

This research used resources of the Advanced Photon Source, a U.S. Department of Energy (DOE) Office of Science User Facility operated for the DOE Office of Science by Argonne National Laboratory under Contract No. DE-AC02-06CH11357. BioCAT was supported by grant P30 GM138395 from the National Institute of General Medical Sciences of the National Institutes of Health. Use of the Pilatus 3 1M detector was provided by grant 1S10OD018090 from NIGMS. The content is solely the responsibility of the authors and does not necessarily reflect the official views of the National Institute of General Medical Sciences or the National Institutes of Health.

## Entry for the Table of Contents

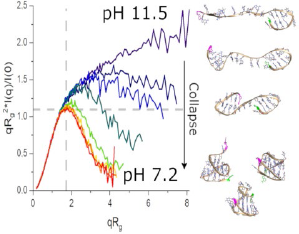

Time-resolved small-angle X-ray scattering (TR-SAXS) has been used to capture the rapid structural collapse of human telomere DNA G-quadruplexes (G4s) using a pH-jump. The monophasic collapse occurs on a time scale of less than 600 ms and precedes much slower folding steps. Our results also reveal that the unfolded ensemble samples transient hairpin conformations prior to a collapse to a molten-globule-like pre-folded particle.

