## Supplemental Material for "Early Events in G-quadruplex Folding Captured by Time-Resolved Small-Angle X-Ray Scattering"

### METHDOS

#### Oligonucleotide Preparation

All oligonucleotides were purchased from Integrated DNA Technologies (IDT, Coralville, IA) as lyophilized, HPLC purified and desalted powders. Stocks solutions of 1 mM were prepared by dilution with MilliQ ultrapure water (18.2 MΩ x cm at 25°C) and stored at -20 °C until use. Oligonucleotide concentrations were determined using the calculated molar extinction coefficients at 260 nm provided by IDT. All samples were further purified in their respective buffer by preparative size-exclusion chromatography (SEC) (Superdex 200 Increase 10/300 GL SEC column, GE Healthcare) prior to concentrating with Pierce protein concentrators (ThermoFisher, #88515). Samples were annealed by diluting into their respective buffer to working concentration followed by heating for 20 min. in 1 L boiling water bath with slow cooling overnight to room temperature. Unless otherwise noted, all experiments were conducted in 8mM potassium phosphate buffer supplemented with 185 mM KCl and pH adjusted to neutral (pH 7.2), alkaline (11.5), or acidic (pH 2.2) using either 1 M KOH or 1 M HCl.

#### Circular Dichroism

Normalized and background-corrected CD spectra were recorded in 1-cm quartz cuvettes at 20°C or 98°C with Jasco J710 or J810 spectropolarimeters adhering to the protocol outlined by Del Villar[1]. Parameters were 220 to 320 or 340 nm wavelength range, 1.0 nm step size, 200 nm/min scan rate, 1.0 nm bandwidth, 2 second integration time, and 4 scan accumulation. Spectra were base-line corrected for background and normalized by strand concentration using the following formula:

$$\Delta\varepsilon = \theta/(32982cl)$$

where $\theta$ is ellipticity in millidegrees, $c$ is molar DNA concentration in mol/L, and $l$ is the path length of the cell in cm. For the pH-jump hand mixing experiments, 2GKU was annealed at 7.1 uM in pH 11.3 phosphate buffer and multiple “pre-jump” scans were measured to ensure it was denatured. At time 0 s, concentrated HCl was added by manual mixing with a pipette such that the pH was rapidly dropped to pH 7.1 (measured after the experiment) with a total dead time of approximately 30 seconds.

#### 1D Nuclear Magnetic Resonance (NMR)

1D $^1$H-NMR spectroscopy was performed on a Bruker Avance Neo 600-MHz instrument equipped with a nitrogen-cooled Prodigy TCI cryoprobe. 2GKU and 2JSL samples were prepared as above, and concentrated to 0.5-1 mM in 250 µL volume using pre-rinsed Pall 3K MWCO concentrators prior to the addition of 5% v/v $D_2O$. Measurements were made at 25°C in standard 3-mm NMR tubes. Minimization of water signal was achieved using a water flip-back pulse sequence.

For the pH-jump experiments, samples were prepared by annealing in alkaline phosphate buffer (pH 11.5) at ~5 µM and concentrated to 2.2 mM in a volume of 500 µL using pre-rinsed Pall 3K MWCO concentrators, followed by addition of 5% v/v $D_2O$. Experiments were performed at 25°C using a 5-mm NMR tube. Folding was initiated by adding 30 µL of a 1M HCl solution to the NMR tube with gentle mixing using a stretched transfer pipette. The approximate dead time between mixing and measurement was 1 min. 45 s. Minimization of water signal was achieved using a water flip-back pulse sequence. 100 measurements were made over 693 seconds with 4 scans averaged per measurement. Measurements were repeatable by adjusting the pH back to 11.5 using the same %v/v addition of 1M KOH and repeating the pH drop to neutral with additional 1 M HCl solution.

**Analytical Ultracentrifugation**

AUC Sedimentation velocity experiments were done using a Beckman Coulter ProteomeLab XL-A analytical ultracentrifuge (Beckman Coulter Inc., Brea, CA) at 20.0°C and using standard 2 sector cells. Each experiment was done at 40k rpm and 150 scans were collected over an 8-hour centrifugation period. Data were analyzed using the program SEDFIT[2] in the continuous c(s) model. Buffer density was determined on a Mettler/Paar Calculating Density Meter DMA 55A at 20.0°C and buffer viscosity was measured on an Anton Paar Automated Microviscometer AMVn. For the calculation of frictional ratio and molecular weight, 0.55 mL/g was used for both single-stranded and G4 partial specific volume[3].

**Equilibrium (EQ)-SAXS**

Size-exclusion chromatography coupled SAXS (SEC-SAXS) was performed at the BioCAT beamline (18ID) at the Advanced Photon Source (APS) at Argonne National Lab. Prepared samples were centrifuged and subsequently loaded onto an equilibrated Superdex 200 Increase 10/300 GL column (Cytiva) maintained at a flow rate 0.7 mL/min (see **Table S1** below) using an AKTA Pure FPLC (GE Healthcare Life Sciences). After passing through the UV monitor, the eluate was directed through the SAXS flow cell, which consists of a 1 mm ID quartz capillary with 20 µm walls. A co-flowing buffer sheath was used to separate the sample and the capillary walls, helping to prevent radiation damage [4]. Scattering intensity was recorded with a Pilatus3 X 1M (Dectris) detector placed 3.628 m from the sample, giving access to a q-range of 0.0044 $Å^{-1}$ to 0.35 $Å^{-1}$. A series of 0.5 second exposures were acquired continuously during elution and the data was reduced using the software *BioXTAS RAW version 2.1.1* [5]. Buffer blanks were created by averaging regions flanking the elution peak and subtracted from exposures selected from the sample elution peak to create the buffer corrected I(q) vs. q curves for subsequent analyses. SAXS sample preparation, data collection, data reduction, analysis, presentation, and interpretation have been done in close accordance with recently published guidelines[6]. All SAXS data have been deposited in the SASBDB (https://www.sasbdb.org/)[7].

**Time-resolved (TR)-SAXS**

Time-resolved SAXS (TR-SAXS) studies were performed at the BioCAT beamline (18ID) at the Advanced Photon Source (APS) at Argonne National Lab. DNA samples were prepared in pH 11.5 buffer as described above and concentrated to 9 mg/mL. The mixing buffer was a matched phosphate buffer at pH 2.2. TR-SAXS studies utilized a 5-inlet laminar flow mixing device[8] capable of measuring events on the 1 – 1500 ms timescale. The central channel contained the DNA samples, the diagonal channels contained the pH 11.5 sample buffer, and the vertical channels contained the pH 2.2 mixing buffer. Hand mixing experiments mimicking the conditions of the TR-SAXS experiment were conducted prior to analysis to ensure that the final pH upon mixing would be ~7.2. Using 2GKU, a concentration series was conducted at 8.7, 4.0, and 2.0 mg/mL to evaluate ideal scattering conditions. It was determined that 4 mg/mL achieved the best S/N and agreement with EQ-SAXS results. Data was collected over ~1 ms to 1.2 s with 118 time points. Reported results are for 2GKU and 2JSL at 4.0 mg/mL.

Standard data reduction methods were used for radial averaging of each measurement. During some of the runs, the first few (time) points were missed due to slight mis-alignment of the sample stream with the beam path and so multiple runs at 4.0 mg/mL were conducted and averaged to obtain the final data sets. This is the reason that the I(0) plots show some sharp discontinuities, as I(0) is related to the concentration of sample. We also note that some data points in the low q range appear to sharply curve up, possibly indicating aggregation. However, orthogonal methods (NMR, AUC) show no detectable aggregation or oligomerization with the 2GKU and 2JSL sequences at these concentrations. The averaged data sets were analyzed using REGALS[9] in the BioXTAS RAW v2.2.2[10] software with the EQ-SAXS data of the neutral and alkaline denatured sequences used to anchor at the beginning and end of the TR-SAXS data sets.

**Molecular dynamics and modeling**

Starting coordinates of the single-stranded 2GKU and 2JSL sequences were generated in UCSF Chimera[11] using the ‘build’ functionality. The PDB structures were then imported into the tleap module of AMBER 2020[12] to generate the sander input files. All simulations were done using the Generalized Born implicit solvation model (igb=8) and equilibrated using sander at 300 K and 1 atm using the following steps: (1) minimization with weak restraints of 10.0 kcal/mol/Å on all nucleic acid residues (2000 cycles of minimization, 500 steepest decent before switching to conjugate gradient) and 16.0 Å cutoff, (2) heating from 0 K to 100 K over 20 ps with 50 kcal/mol/Å restraints on all nucleic acid residues, (3) minimization of the system without restraints (2500 cycles, 1000 steepest decent before switching to conjugate gradient) with 16 Å cutoff, (4) heating from 100 K to 300 K over 20 ps with weak restraints of 10.0 kcal/mol/Å on nucleic acid residues, and (5) equilibration at 1 atm for 100 ps with weak restraints of 10.0 kcal/mol/Å on nucleic acids. The resulting coordinate files from equilibration were then used as input for duplicate 500 ns of unrestrained, solvated MD simulations using pmemd with GPU acceleration in the isothermal isobaric ensemble (P = 1 atm, T = 300 K) with the DNA OL15 force field. Random initial velocities were achieved using the date and time of the simulation (ig=-1). 2.0 fs time steps were used with bonds involving hydrogen frozen using SHAKE (ntc = 2). Trajectories were analyzed using the CPPTRAJ module in the AmberTools20[12] package.

Molecular model ensembles were derived using the Ensemble Optimization Method 2.1[13] program from the ATSAS suite of tools. A total of 10,000 snapshots equally spaced across the two 500 ns combined trajectories for both 2GKU and 2JSL were used to generate models that were included in each EOM pool (i.e., a total of 10,000 models made up each respective pool). GAJOE was used in pool "-p" mode, with maximum curves per ensemble set to 20, minimum curves per ensemble set to 5, constant subtraction allowed, curve repetition allowed, and the genetic algorithm (GA) repeated 100 times. In brief, EOM takes a large pool of macromolecules covering as much conformational space as possible (and reasonable) and selects from this pool a sub-ensemble of conformers that recapitulate the experimental scattering. The ensemble is the subset of weighted theoretical curves from conformations that minimizes the discrepancy $\chi^2$:

$$\chi^2 = \frac{1}{K-1}\sum_{j=1}^{K}\left[\frac{\mu I(s_j) - I_{exp}(s_j)}{\sigma(s_j)}\right]^2$$

where $I_{exp}(s_j)$ is the experimental scattering, $I(s_j)$ is the calculated scattering, $K$ is the number of experimental points, $\sigma(s_j)$ are standard deviations, and $\mu$ is a scaling factor[14]. Ten independent EOM runs were performed for each system to ensure that convergance was achieved.

Calculations of 5' and 3' distances were performed in UCSF Chimera[11] using the 'distance' command. SASA values were determined using the program FreeSASA[15] using default atom radii, slices (20), and a probe size of 1.400 Å.

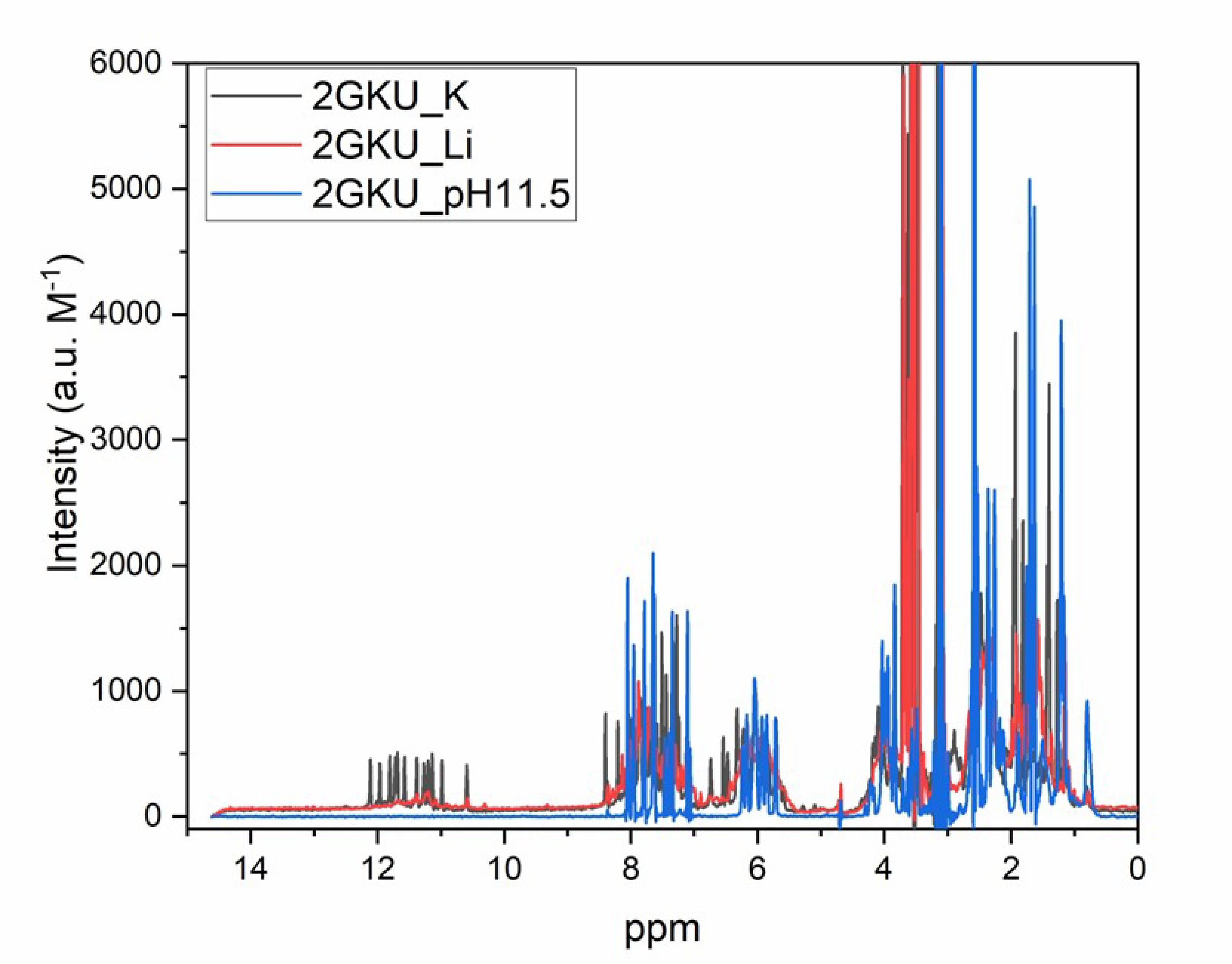


**Figure S1.** NMR spectra of 2GKU under various solution conditions. The alkaline denatured and $Li^+$ conditions show a sharp reduction in the G4 imino shifts in the range of 10-12 ppm.

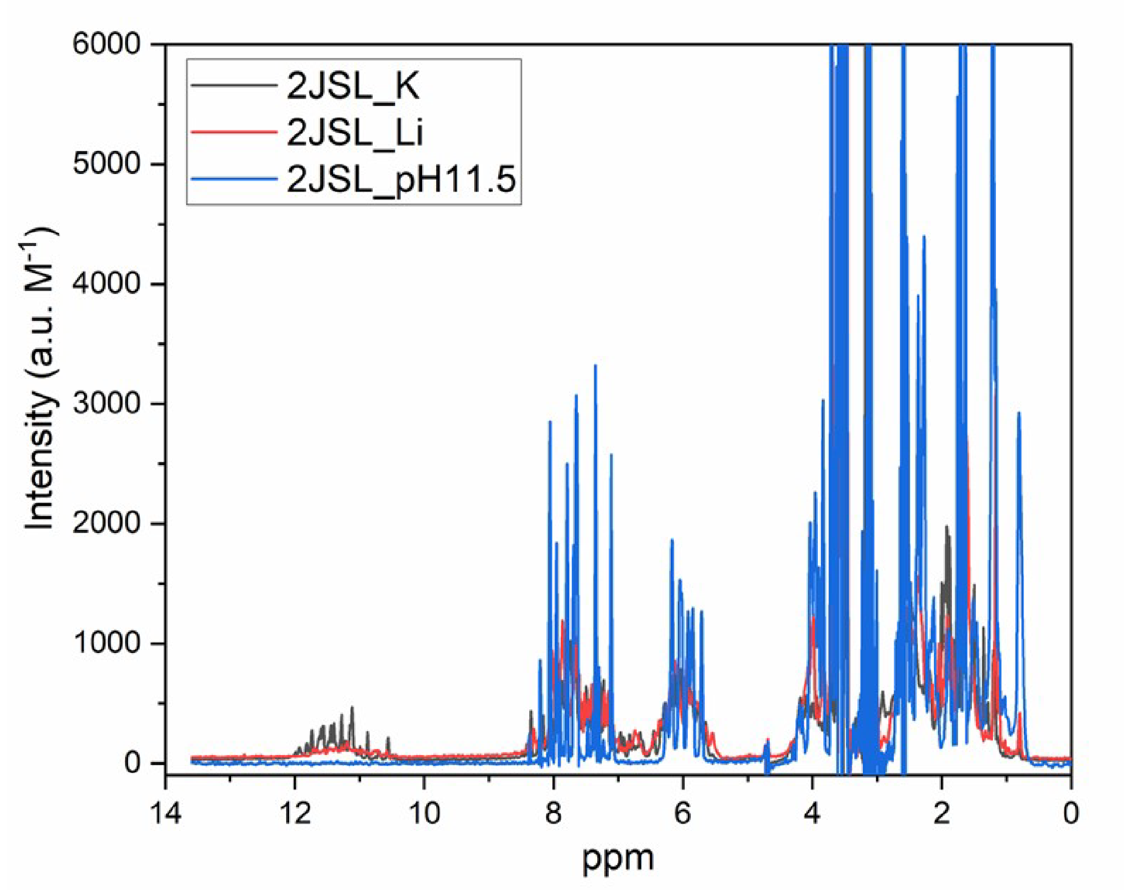


**Figure S2.** NMR spectra of 2JSL under various solution conditions. The alkaline denatured and $Li^+$ conditions show a sharp reduction in the G4 imino shifts in the range of 10-12 ppm.

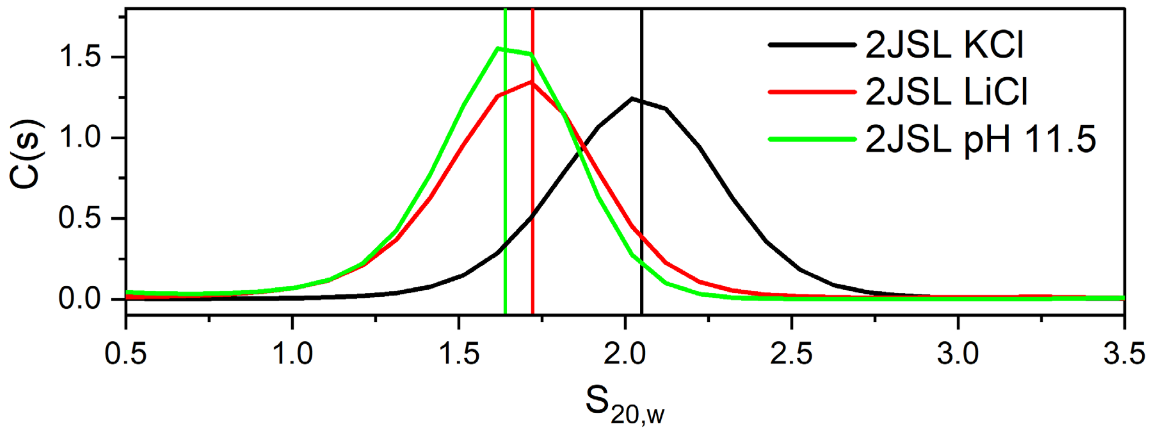


**Figure S3.** AUC-SV analysis of 2JSL under various buffer conditions showing a reduction in sedimentation coefficient in alkaline or LiCl conditions.

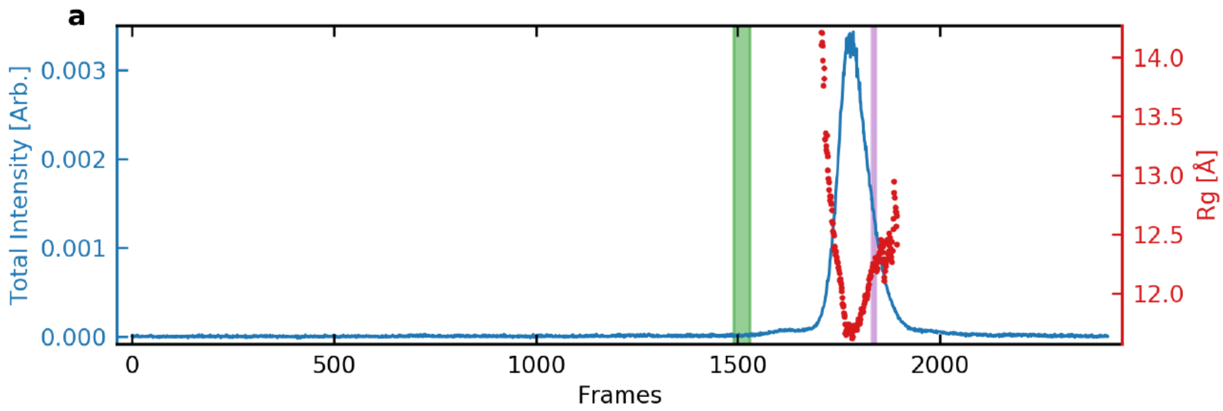


**Figure S4.** SEC-SAXS elution profile for 2GKU at pH 7.2 with KCl. Plot shows the series intensity (blue, left axis) vs. frame and $R_g$ vs. frame (red, right axis). The green shaded region depicts the buffer region and purple shaded region shows the sample region used in subsequent analyses.

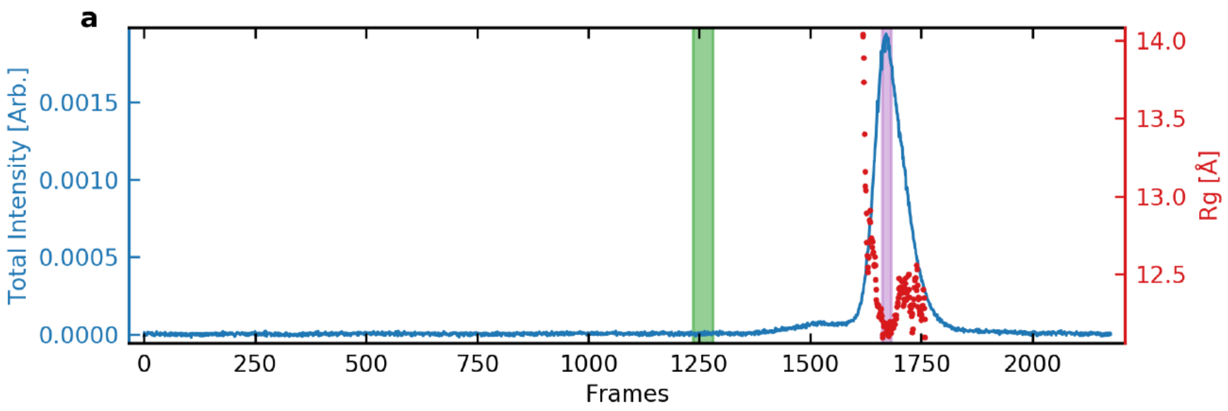


**Figure S5.** SEC-SAXS elution profile for 2JSL at pH 7.2 with KCl. Plot shows the series intensity (blue, left axis) vs. frame and $R_g$ vs. frame (red, right axis). The green shaded region depicts the buffer region and purple shaded region shows the sample region used in subsequent analyses.

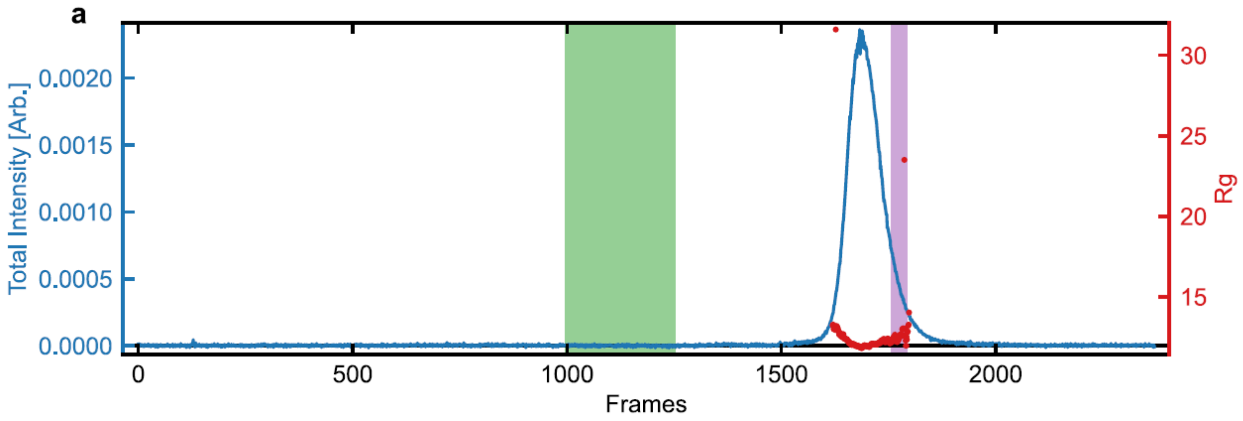


**Figure S6.** SEC-SAXS elution profile for 2GKU at pH 7.2 with LiCl. Plot shows the series intensity (blue, left axis) vs. frame and $R_g$ vs. frame (red, right axis). The green shaded region depicts the buffer region and purple shaded region shows the sample region used in subsequent analyses.

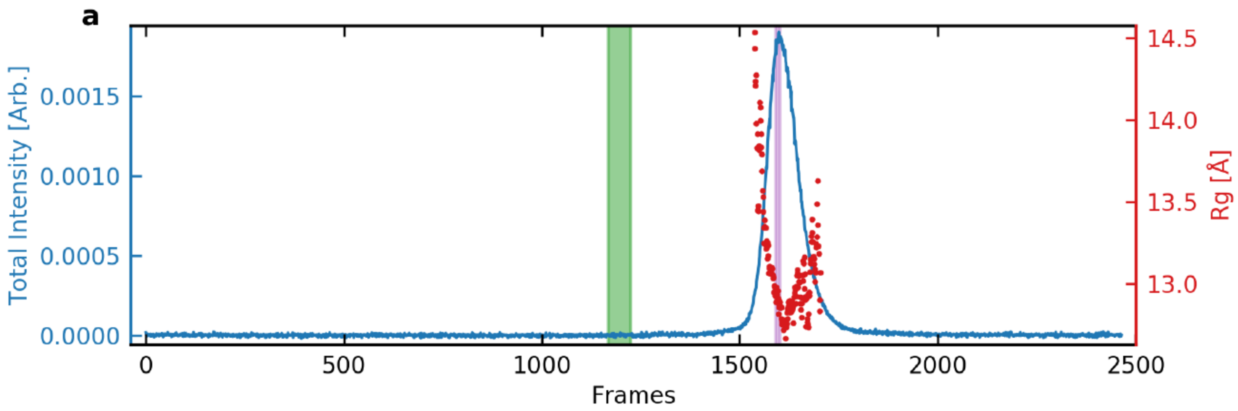


**Figure S7.** SEC-SAXS elution profile for 2JSL at pH 7.2 with LiCl. Plot shows the series intensity (blue, left axis) vs. frame and $R_g$ vs. frame (red, right axis). The green shaded region depicts the buffer region and purple shaded region shows the sample region used in subsequent analyses.

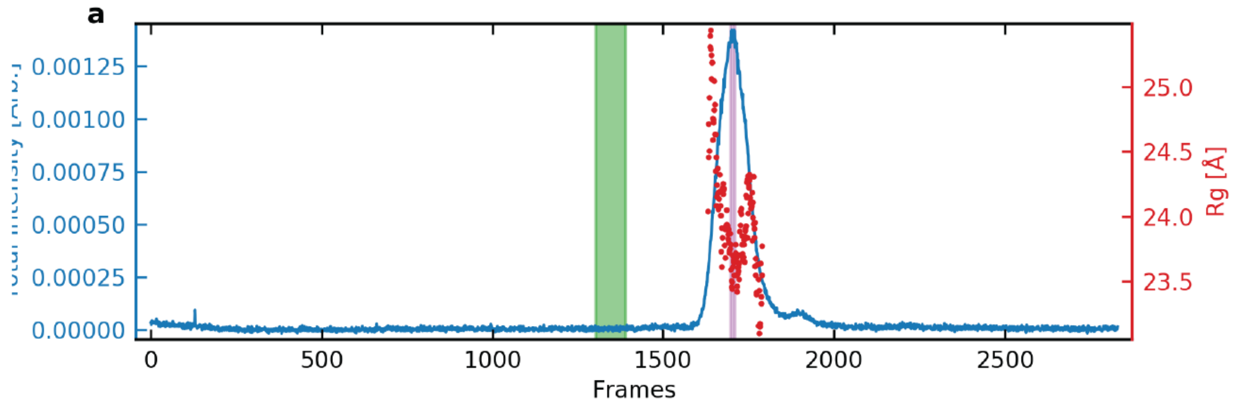


**Figure S8.** SEC-SAXS elution profile for 2GKU at pH 11.5 with KCl. Plot shows the series intensity (blue, left axis) vs. frame and $R_g$ vs. frame (red, right axis). The green shaded region depicts the buffer region and purple shaded region shows the sample region used in subsequent analyses.

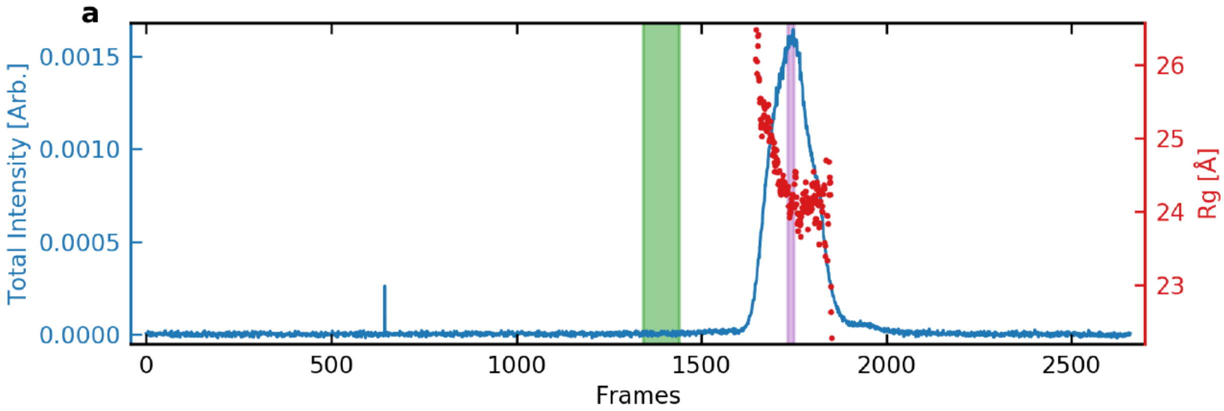


**Figure S9.** SEC-SAXS elution profile for 2JSL at pH 11.5 with KCl. Plot shows the series intensity (blue, left axis) vs. frame and $R_g$ vs. frame (red, right axis). The green shaded region depicts the buffer region and purple shaded region shows the sample region used in subsequent analyses.

(a) Sample Details.

| | 2GKU pH 7.2 (KCl) | 2JSL pH 7.2 (KCl) | 2GKU pH 7.2 (LiCl) | 2JSL pH 7.2 (LiCl) | 2GKU pH 11.5 (KCl) | 2JSL pH 11.5 (KCl) |
|---|---|---|---|---|---|---|
| Organism | synthetic | synthetic | synthetic | synthetic | synthetic | synthetic |
| Source | IDT | IDT | IDT | IDT | IDT | IDT |
| Extinction coefficient (nearest neighbor approximation) ($M^{-1}$ $cm^{-1}$) | 244300 | 253100 | 244300 | 253100 | 244300 | 253100 |
| M from chemical composition (Da) | 7575 | 7879 | 7575 | 7879 | 7575 | 7879 |
| SEC-SAXS column, 10 x 300 Superdex 75 | | | | | | |
| Loading concentration (mg/mL) | 10.3 | 21.1 | 15 | 12.5 | 12.2 | 19.9 |
| Injection volume (µL) | 330 | 113 | 128 | 263 | 233 | 137 |
| Flow rate (mL/min) | 0.7 | 0.7 | 0.7 | 0.7 | 0.7 | 0.7 |
| Solvent (solvent blanks taken from SEC flow through prior to elution of protein) | 8 mM $PO_4^{2-}$, 185 mM KCl, 1 mM EDTA, pH 7.2 | 8 mM $PO_4^{2-}$, 185 mM KCl, 1 mM EDTA, pH 7.2 | 8 mM $PO_4^{2-}$, 185 mM LiCl, 1 mM EDTA, pH 7.2 | 8 mM $PO_4^{2-}$, 185 mM LiCl, 1 mM EDTA, pH 7.2 | 8 mM $PO_4^{2-}$, 185 mM KCl, 1 mM EDTA, pH 11.5 | 8 mM $PO_4^{2-}$, 185 mM KCl, 1 mM EDTA, pH 11.5 |

(b) SAXS data-collection parameters.

| | |
|---|---|
| Instrument/data processing | BioCAT facility at the Advanced Photon Source beamline 18ID with Pilatus3 1M (Dectris) detector |
| Wavelength (Å) | 1.033 |
| Beam size (µm) | 150 (h) x 25 (v) |
| Camera length (m) | 3.655 |
| q measurement range ($Å^{-1}$) | 0.0044-0.35 |
| Absolute scaling method | N/A |

| | | | | | | |
|---|---|---|---|---|---|---|
| Normalization | To incident intensity, by ion chamber counter | | | | | |
| Monitoring for radiation damage | Automated frame-by-frame comparison of relevant regions | | | | | |
| Exposure time, number of exposures | 0.5 s exposure time with a 2s total exposure period (0.5 s on, 1.5 s off) of entire SEC elution | | | | | |
| Sample configuration | SEC-SAXS. Size separation by an AKTA Pure with a Superdex 75 Increase 10/300 GL column. SAXS data measured in a 1.5 mm ID quartz capillary | | | | | |
| Sample temperature (°C) | 22 | | | | | |
| (c) Software employed for SAXS data reduction, analysis, and interpretation. | | | | | | |
| SAXS data reduction | Radial averaging; frame comparison, averaging, and subtraction done using BioXTAS RAW 2.1.1[16] | | | | | |
| Extinction coefficient estimate | Nearest neighbor approximation | | | | | |
| Basic analyses: Guinier, P(r), $V_p$ | Guinier fit, Kratky analysis, and molecular weight using BioXTAS RAW 2.1.1, P(r) function using PRIMUSqt (ATSAS v2.8.4[17]) | | | | | |
| Shape/bead modelling | N/A | | | | | |
| Atomic structure modelling | UCSF Chimera v1.11 & AMBER 2020 | | | | | |
| Three-dimensional graphic model representations | UCSF Chimera v1.11 | | | | | |
| (d) Structural parameters. | | | | | | |
| Guinier analysis | 2GKU pH 7.2 (KCl) | 2JSL pH 7.2 (KCl) | 2GKU pH 7.2 (LiCl) | 2JSL pH 7.2 (LiCl) | 2GKU pH 11.5 (KCl) | 2JSL pH 11.5 (KCl) |

| | | | | | | |
|---|---|---|---|---|---|---|
| I(0) ($cm^{-1}$) | 0.0112 ± 0.00002 | 0.0153 ± 0.00002 | 0.0166 ± 0.000008 | 0.0152 ± 0.00002 | 0.0178 ± 0.00005 | 0.0206 ± 0.00003 |
| $R_g$ (Å) | 12.27 ± 0.04 | 12.13 ± 0.03 | 11.92 ± 0.01 | 12.89 ± 0.04 | 24.26 ± 0.17 | 24.26 ± 0.11 |
| qmin ($Å^{-1}$) | 0.00586 | 0.00729 | 0.011 | 0.00872 | 0.00643 | 0.00843 |
| $qR_g$ max | 1.2576 | 1.146 | 1.3099 | 1.2066 | 0.9949 | 1.0365 |
| Coefficient of correlation, $R^2$ | 0.9796 | 0.9883 | 0.9988 | 0.9892 | 0.9776 | 0.9902 |
| SAXS MW (Ratio to Expected) (kDa) | 9.7 (1.28) | 9.9 (1.26) | 9.0 (1.19) | 9.5 (1.21) | 15.1 (1.99) | 15.7 (1.99) |
| P(r) analysis (GNOM) | | | | | | |
| I(0) ($cm^{-1}$) | 0.0112 ± 0.00002 | 0.0154 ± 0.00002 | 0.00167 ± 0.0000103 | 0.0152 ± 0.00002 | 0.0178 ± 0.00004 | 0.0207 ± 0.00003 |
| Rg (Å) | 12.23 ± 0.03 | 12.20 ± 0.03 | 11.97 ± 0.02 | 13.18 ± 0.10 | 24.56 ± 0.11 | 25.03 ± 0.08 |
| $D_{max}$ (Å) | 38 | 39 | 43 | 51 | 83 | 87 |
| $\chi^2$ | 1.201 | 1.354 | 1.070 | 1.022 | 1.155 | 1.058 |
| Porod volume ($Å^{-3}$) (ratio $V_p$/calculated M) | 4500 | 4670 | 3920 | 4540 | 12400 | 12500 |

(e) Shape model-fitting results

| | 2GKU pH 7.2 (KCl) | 2JSL pH 7.2 (KCl) | 2GKU pH 7.2 (LiCl) | 2JSL pH 7.2 (LiCl) | 2GKU pH 11.5 (KCl) | 2JSL pH 11.5 (KCl) |
|---|---|---|---|---|---|---|
| (f) Atomistic modelling. | | | | | | |
| Crystal structures/atomic coordinate files | | | | | Modeled | Modeled |
| q range for modelling | | | | | 0.00643-0.3 | 0.00843-0.3 |
| EOM GAJOE 2.1 (min ensembles = 5, max = 20, default parameters) | | | | | | |
| $\chi^2$ | | | | | 1.162 | 1.126 |
| Rflex (random) / Rsigma | | | | | 84.25 (90.67) / 0.85 | 73.58 (87.66) / 0.63 |

| | | | | | | |
|---|---|---|---|---|---|---|
| Constant subtracted | | | | | 0 | 0 |
| No. of representative structures | | | | | 9 | 3 |
| Final ensemble $R_g$ (Å), $D_{max}$ (Å) | | | | | 24.56, 76.94 | 25.54, 79.24 |

(g) SASBDB IDs for data and models.

| | | | | | | |
|---|---|---|---|---|---|---|
| ID | TBD | TBD | TBD | TBD | TBD | TBD |

**Table S1.** Tabulated equilibrium SEC-SAXS data acquisition, reduction, analysis, results and SASBDB identifiers.

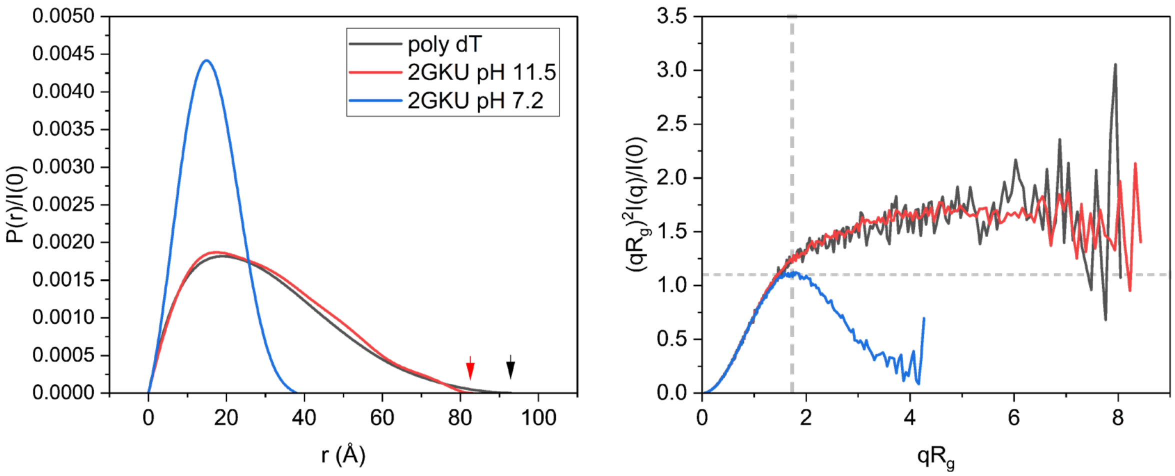


**Figure S10.** Normalized P(r) and Dimensionless Kratky plots comparing folded 2GKU in KCl at pH 7.2 (blue), pH 11.5 (red), and poly dT$_{24}$ (black) at pH 7.2 with KCl.

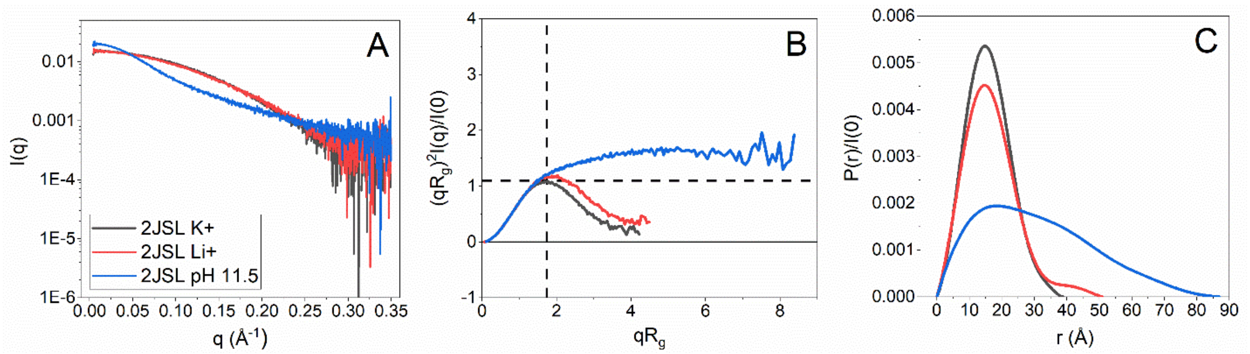


**Figure S11.** (A) Scattering profiles, (B) Dimensionless Kratky plots, and (C) Normalized P(r) distributions of 2JSL in KCl at pH 7.2 (black), LiCl at pH 7.2 (red), and KCl at pH 11.5 (blue).

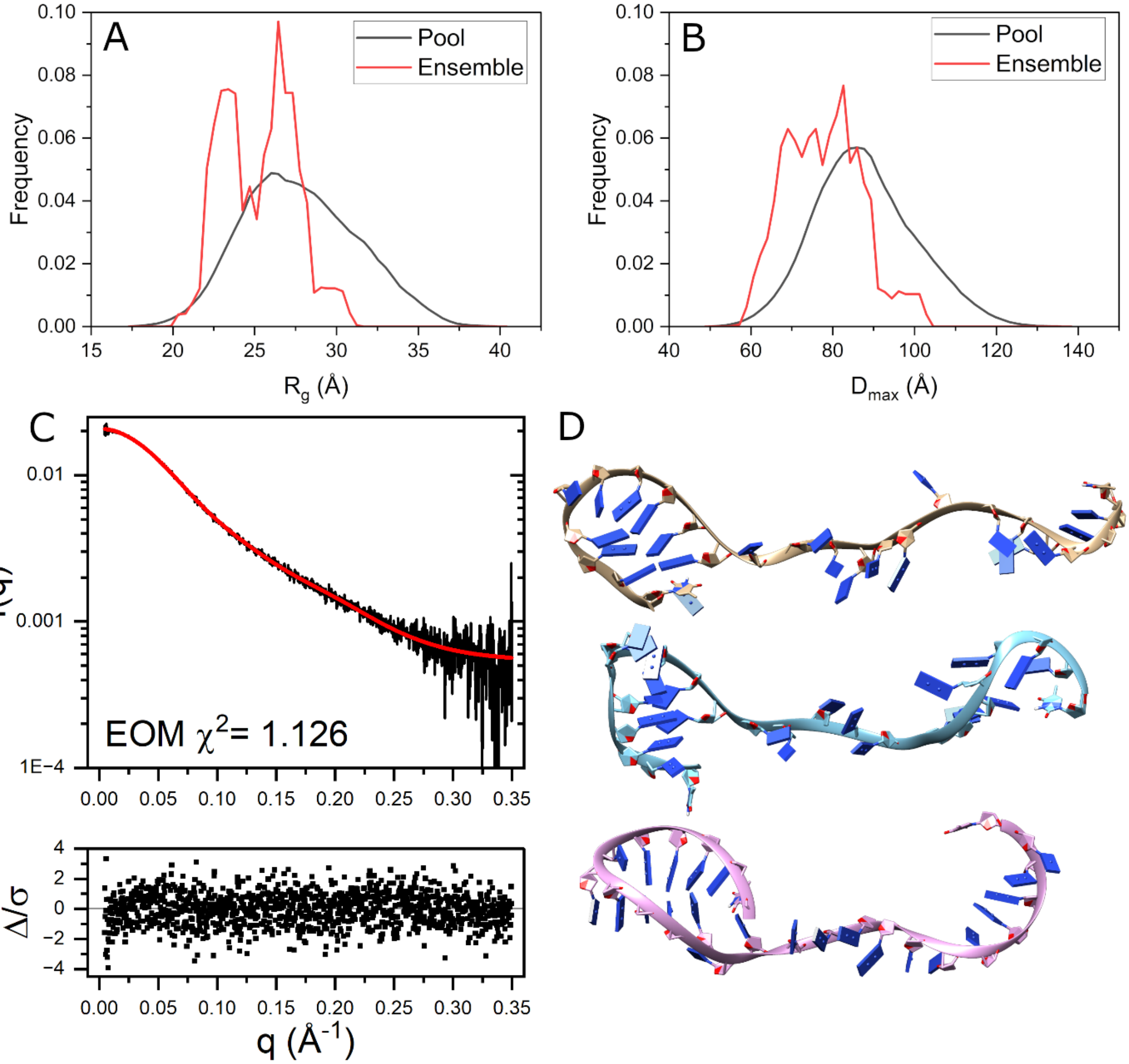


**Figure S12.** EOM analysis results for 2JSL. (A-B) EOM distributions for the radius of gyration (A) and $D_{max}$ (B) for the total pool of conformers (black) and the selected ensemble of flexible structures (red). (C) pH 11.5 scattering curve with EOM fit overlaid in red and residuals below. (D) Best fit ensemble of conformers chosen by EOM from duplicate 500 ns implicitly solvated MD simulations starting from single-stranded 2JSL showing the most extended (top) to the most compact conformations (bottom) oriented with 5' end on the left. Calculated $D_{max}$ values and % ensemble weight are given above each model. EOM statistics: $R_{flex}$ (random) / $R_{sigma}$: ~ 73.58% (~ 87.66%) / 0.63.

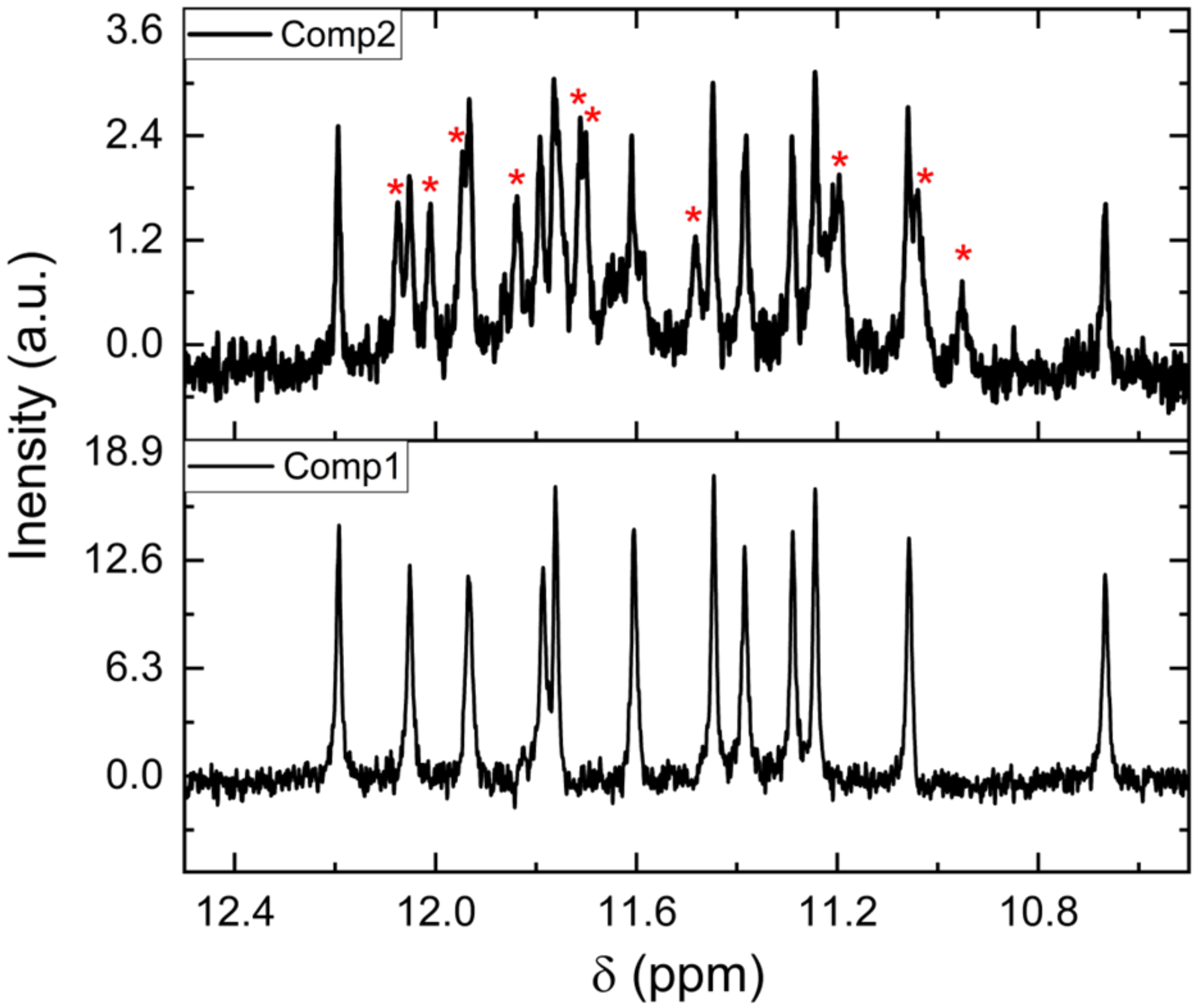


**Figure S13.** SVD results of hand-mixing pH jump 1D-NMR experiments showing the imino proton regions of the two major spectral components. Red asterisks indicate imino proton shifts that disappear rapidly after pH jump and may correspond to the early spectral intermediate observed by CD hand mixing in main text Figure 4. Component 1 is consistent with the imino shifts reported for the folded hybrid-1 2GKU.

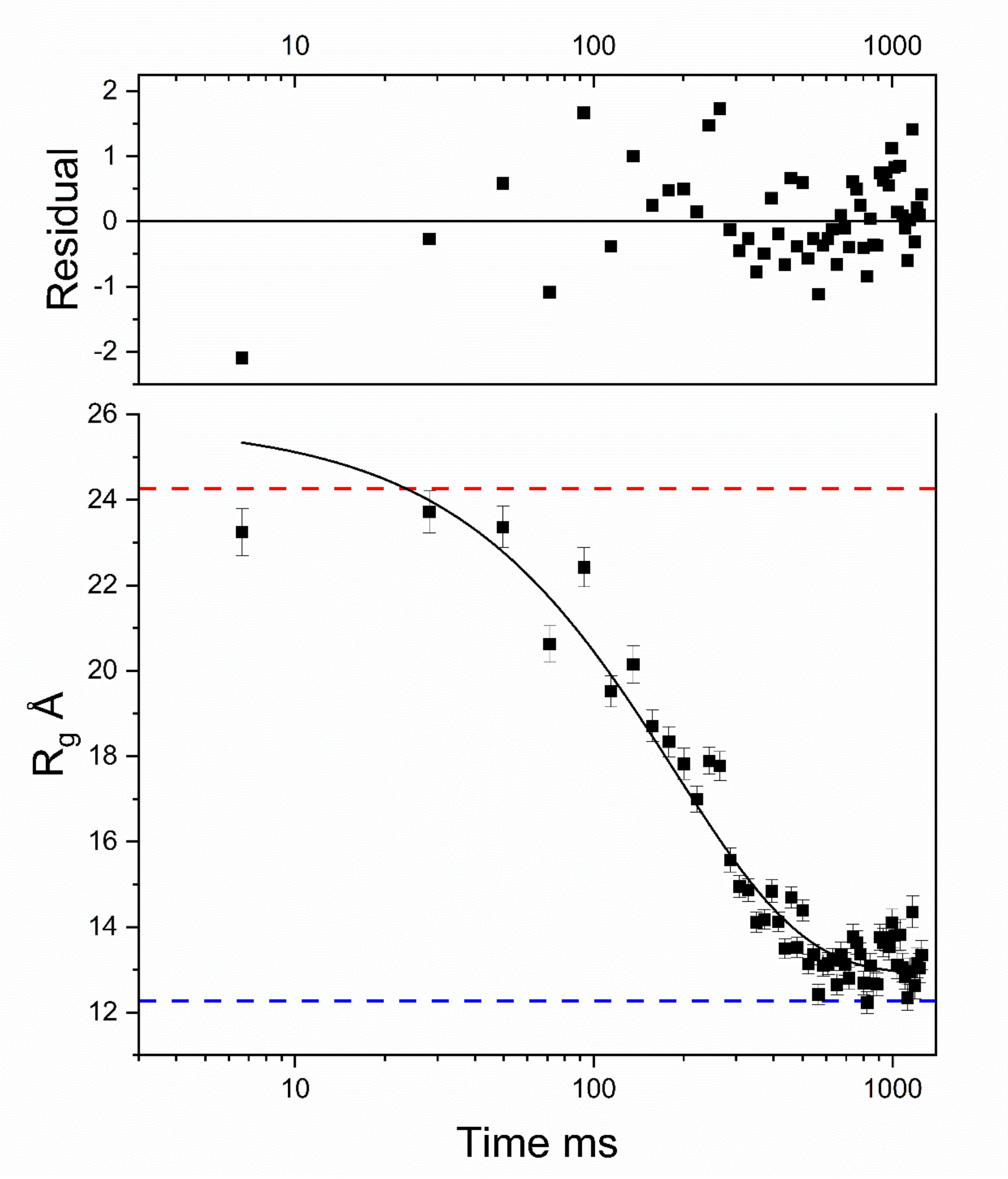


**Figure S14.** Time course of $R_g$ changes for 2GKU following an 11.5 to 7.2 pH jump. Experimental data (square symbols with error bars) are shown over the range of 0 – 1200 milliseconds. The solid line is the best least-squares fit to an exponential decay with a time constant of 187 ± 13 milliseconds. The top panel shows the residuals of the fit. Red and blue dashed lines indicate the EQ-SAXS measured $R_g$ values for the alkaline denatured and pH 7.2 equilibrated 2GKU, respectively.

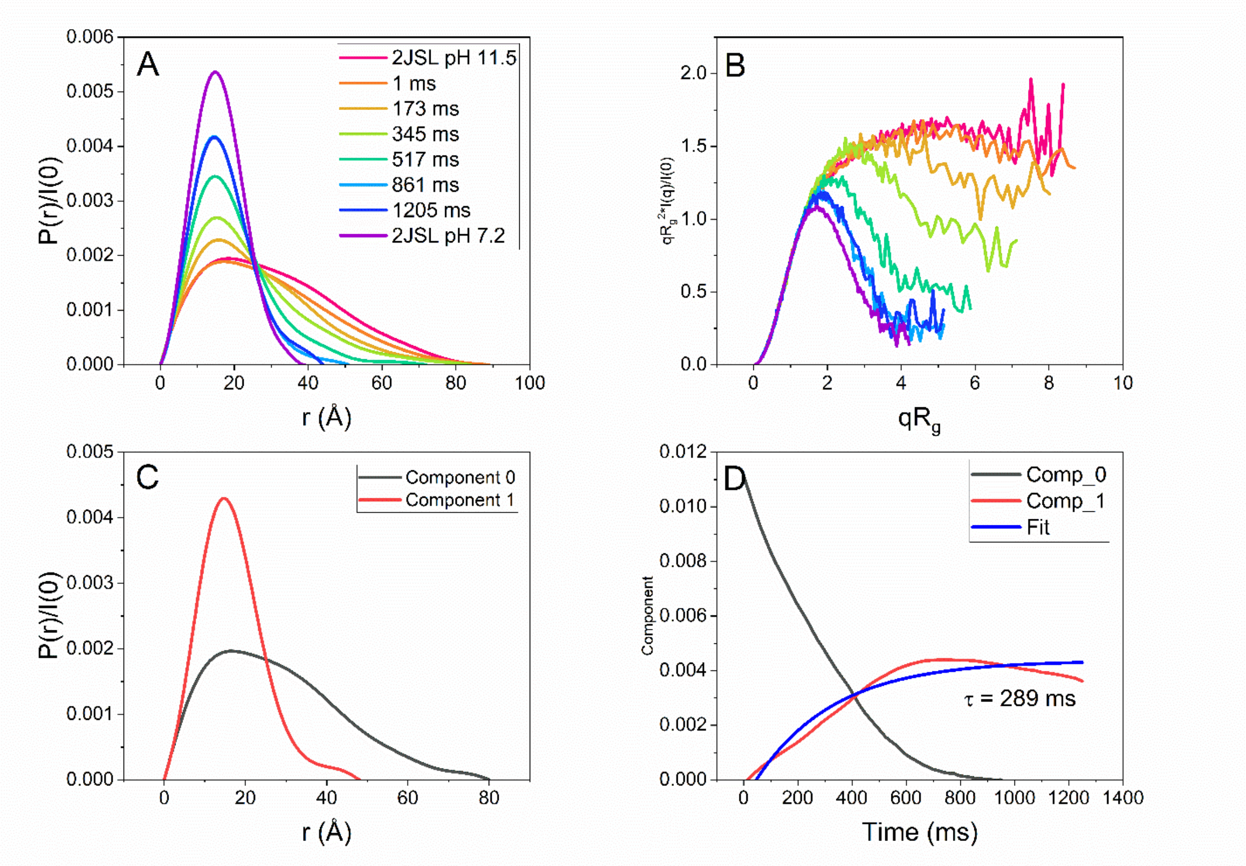


**Figure S15.** Time-resolved SAXS results of 2JSL following pH jump from 11.5 to 7.2. (A) Selected normalized P(r) distributions of the pH-induced structural collapse of 2GKU bracketed by the equilibrium SAXS profiles showing the conversion from an extended unstructured species to a globular and compact particle of nearly identical size and shape as the folded hybrid 1 form. (B) Dimensionless Kratky plots (data re-binned for clarity with log mode and re-bin factor 4) showing the transition from the denatured flexible chain to a compact globular form that is nearly identical to the equilibrium 2GKU scattering. (C) REGALS derived regularized P(r) distributions comparing the two deconvoluted components. (D) Regularized component concentration profiles from REGALS deconvolution with single exponential decay relaxation times overlaid and fit shown in blue. Red and black curves correspond to the solid red and black P(r) distributions in C.

[12] I. Y. B.-S. D.A. Case, S.R. Brozell, D.S. Cerutti, T.E. Cheatham, III, V.W.D. Cruzeiro, T.A. Darden, R.E. Duke, D. Ghoreishi, M.K. Gilson, H. Gohlke, A.W. Goetz, D. Greene, R Harris, N. Homeyer, S. Izadi, A. Kovalenko, T. Kurtzman, T.S. Lee, S. LeGrand, P. Li, C. Lin, J. Liu, T. Luchko, R. Luo, D.J. Mermelstein, K.M. Merz, Y. Miao, G. Monard, C. Nguyen, H. Nguyen, I. Omelyan, A. Onufriev, F. Pan, R. Qi, D.R. Roe, A. Roitberg, C. Sagui, S. Schott-Verdugo, J. Shen, C.L. Simmerling, J. Smith, R. Salomon-Ferrer, J. Swails, R.C. Walker, J. Wang, H. Wei, R.M. Wolf, X. Wu, L. Xiao, D.M. York and P.A. Kollman, **2018**.
